# JAK-STAT Signaling Enables Lineage Plasticity-driven AR Targeted Therapy Resistance

**DOI:** 10.1101/2021.11.02.466956

**Authors:** Su Deng, Choushi Wang, Yunguan Wang, Yaru Xu, Xiaoling Li, Nickolas A Johnson, U-Ging Lo, Lingfan Xu, Julisa Gonzalez, Lauren A Metang, Jianfeng Ye, Carla Rodriguez Tirado, Kathia Rodarte, Zhiqun Xie, Carlos Arana, Valli Annamalai, Jer-Tsong Hsieh, Donald J. Vander Griend, Bo Li, Tao Wang, Ping Mu

## Abstract

Emerging evidence indicates that various cancers can gain resistance to targeted therapies by acquiring lineage plasticity. Although various genomic and transcriptomic aberrations correlate with lineage plasticity-driven resistance, the molecular mechanisms of acquiring lineage plasticity have not been fully elucidated. Through integrated transcriptomic and single cell RNA-Seq (scRNA-Seq) analysis of more than 80,000 cells, we reveal for the first time that the Janus kinase (JAK)-signal transducer and activator of transcription (STAT) signaling is a crucial executor in promoting lineage plasticity-driven AR targeted therapy resistance in prostate cancer. Ectopic activation of JAK-STAT signaling is specifically required for the AR targeted therapy resistance of subclones expressing multilineage, stem-like and epithelial-to-mesenchymal transition (EMT) lineage transcriptional programs and represents a potential therapeutic target for overcoming AR targeted therapy resistance.

**One-Sentence Summary:** JAK-STAT signaling is a crucial executor in promoting lineage plasticity-driven AR therapy resistance in prostate cancer.

## Main Text

Despite the clinical success of targeted therapies directed towards specific driver oncogenes in many cancers, resistance to these therapies often emerges quickly, resulting in poor clinical outcomes. Metastatic castration resistant prostate cancer (mCRPC) serves as a salient example of this phenomenon, whereby resistance to the Androgen Receptor (AR) targeted therapies, such as enzalutamide, with subsequent disease progression occurs rapidly and is often inevitable (*1–3*). Several mechanisms have been revealed to confer resistance to AR targeted therapy through either the restoration of AR signaling (*4*) or the bypass of AR signaling via other transcription factors (*5–7*). Recently, emerging evidence has demonstrated a third mechanism of resistance called lineage plasticity, whereby the luminal prostate epithelial cells transition to a lineage plastic state independent of AR(*8*). The acquisition of lineage plasticity may result in cells transitioning to a multi-lineage, stem cell-like and lineage plastic state followed by redifferentiation to new lineages, or possibly direct trans-differentiation to a different lineage, such as the lineage with neuroendocrine (NE) differentiation (*8*).

One example of the lineage plasticity-driven resistance occurs in mCRPC with concurrent loss-of-function of TP53 and RB1, which is then accompanied by ectopic activation of SOX2(*9, 10*). Similar cases of lineage plasticity have been observed in mCRPC carrying various genomic and transcriptional aberrations, including but not limited to aberrations in PTEN, FOXA1, BRN2, SOX11, N-MYC, PEG10, CHD1, REST, and BRG1 (*7, 11–18*). This lineage plasticity-driven resistance in mCRPC parallels examples documented in EGFR-mutant lung adenocarcinoma, ER-positive breast cancers, and BRAF-mutant melanoma (*19–22*). However, the molecular mechanism that promotes lineage plasticity in many mCRPC subtypes, especially in the context of TP53/RB1-deficiency, is not fully understood. Furthermore, therapeutic approaches targeting lineage plasticity-driven resistance are not currently available, underlying the unmet clinical urgency to identify druggable targets that drive lineage plasticity.

In this study, through a multi-disciplinary approach integrating 3D organoid modeling, as well as bulk and single cell RNA-Seq (scRNA-Seq) analysis, we reveal that the ectopic activation of JAK-STAT signaling pathway is required for the lineage plasticity-driven AR targeted therapy resistance in mCRPC with TP53/RB1-deficiency and SOX2 upregulation. For the first time, our scRNA-Seq results revealed that JAK-STAT signaling is specifically required for the AR therapy resistance of subclones expressing multilineage, stem-like and lineage plastic survival transcriptional programs, but not the subclones only expressing the NE-like lineage program. We also demonstrate that both genetic and pharmaceutical inactivation of key components of the JAK-STAT signaling pathway, including JAK1 and STAT1, re-sensitize the resistant mCRPC tumors to AR targeted therapy. Collectively, these findings suggest that the upregulation of JAK-STAT signaling pathway is a crucial executor driving lineage plasticity, which enables us to identify potential therapeutic targets to overcome AR targeted therapy resistance.

To investigate the underlying molecular driver of lineage plasticity and resistance in the TP53/RB1-deficient mCRPC, we first inquired which transcriptional programs changed concomitantly with the loss of TP53 and RB1, as well as the upregulation of SOX2. By leveraging a series of LNCaP/AR cell lines we have previously generated(*10*), we initially profiled the transcriptomic changes induced by TP53/RB1-deficiency and overexpression of SOX2 in four cell lines which were not exposed to the AR therapy drug enzalutamide, specifically the shNT, shTP53/RB1, shTP53/RB1/SOX2, and SOX2-OE (overexpression) cell lines. As expected, these genetic modifications led to global transcriptomic changes, and Gene Set Enrichment Analysis (GSEA) revealed significantly altered pathways (Fig.1A, fig.S1A). Notably, the JAK-STAT signaling pathway is among the most significantly upregulated pathways altered by TP53/RB1-loss and SOX2 overexpression, while, in contrast, it is downregulated in TP53/RB1/SOX2 triple knockdown cells (Fig.1A, fig.S1B-D). To decipher how these transcriptional changes specifically contribute to the AR therapy resistance, we continued to investigate pathway aberrations by profiling a second set of cell lines and examined the signaling pathway alterations upon enzalutamide treatment in comparison to vehicle (shNT-Enz/Veh, shTP53/RB1-Enz/Veh). Again, we uncovered that the components of the JAK-STAT signaling pathway were consistently upregulated in the resistant cells treated with enzalutamide (fig.S1A, E, F). Interestingly, the JAK-STAT pathway did not significantly change in shNT cells treated with enzalutamide, suggesting it has a specific role in the context of TP53/RB1 deficiency (fig.S1A).

**Fig. 1:**
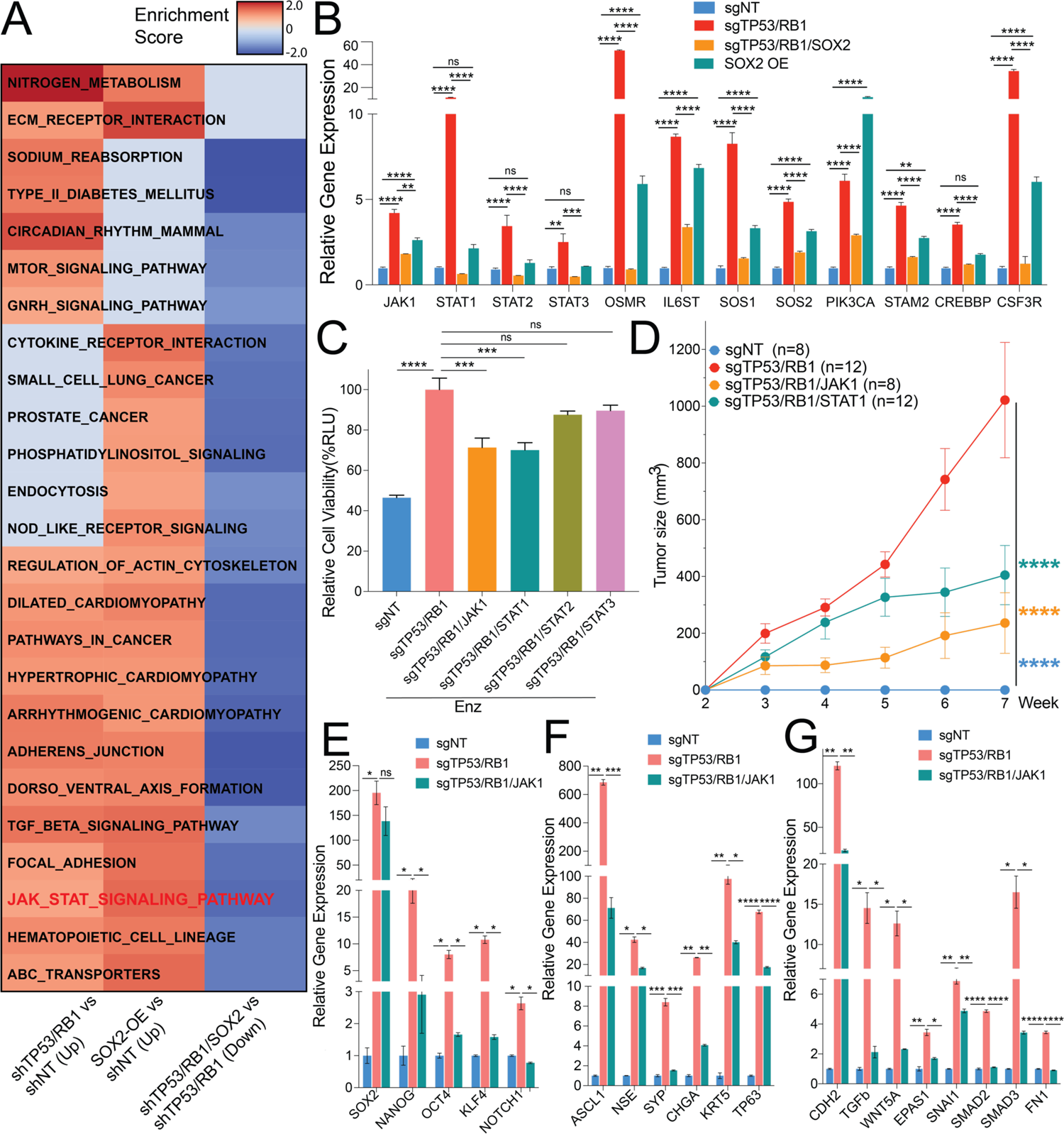
JAK-STAT signaling is required for lineage plasticity and enzalutamide resistance in TP53/RB1-deficient mCRPC. (**A**) Heatmap represents the significantly changed signaling pathways in LNCaP/AR cell lines transduced with annotated shRNAs based on GSEA analysis. Three comparations are presented. Reads from 3 biological replicates in each group were used for analysis. (**B**) Relative gene expression of canonical genes being activated in the JAK-STAT signaling pathway in LNCaP/AR cells transduced with annotated guide RNAs. (**C**) Relative cell number of LNCaP/AR cells transduced with annotated CRISPR guide RNAs. Cells were treated with 10 µM enzalutamide (Enz) for 8 days and cell numbers (viability) were measured using CellTiter-Glo assay, all normalized to sgTP53/RB1 group. p values were calculated using one-way ANOVA. (**D**) Tumor growth curve of xenografted LNCaP/AR cells transduced with annotated guide RNAs in castrated mice. Enz denotes enzalutamide treatment at 10 mg/kg from day 1 of grafting. Veh denotes 0.5% CMC+0.1% Tween 80. p values were calculated using two-way ANOVA. (**E**) Relative expression of canonical stem cell-like lineage marker genes in LNCaP/AR cells transduced with annotated guide RNAs. (F) Relative expression of canonical basal and NE cell-like lineage marker genes in LNCaP/AR cells transduced with annotated guide RNAs. (**G**) Relative expression of canonical EMT lineage marker genes in LNCaP/AR mCRPC cells transduced with annotated guide RNAs. For all panels unless otherwise noted, mean ± s.e.m. is represented and p values were calculated using two-way ANOVA. **** p<0.0001. *** p<0.001. ** p<0.01. * p<0.05. See also **fig.S1-7** and **table S1**.

In normal tissues, the JAK-STAT signaling pathway regulates various biological processes, including immune response, inflammation, embryonic development, cell fate decision, differentiation, and hematopoiesis(*23, 24*). Notably, numerous lines of evidence from both mammalian and *Drosophila* systems implicate that JAK-STAT signaling regulates stem cell self-renewal and multi-lineage differentiation(*23, 25*), indicating its potential role in regulating cellular lineage plasticity. The biological consequence of JAK-STAT activation on tumorigenesis is complicated and considered a “double-edged sword.” On the one hand, JAK-STAT signaling promotes antitumor immune surveillance and is associated with a favorable clinical outcome in some cancers, including colorectal cancer and head and neck squamous cell carcinoma(*23*). Conversely, the constitutive activation of JAK-STAT signaling has been correlated with poor clinical outcomes in hematological malignancies and many solid tumors, including melanoma, glioblastoma, head, neck, lung, pancreas, breast, rectal, and prostate cancers (PCa)(*23, 26–28*). This “double-edged sword” effect of JAK-STAT activation is particularly puzzling in the case of IL-6/STAT3, as IL6-induced STAT3 has been reported to promote PCa NE-differentiation and cell cycle arrest(*29–31*), while antagonizing the AR inhibition induced PCa cell apoptosis and proliferation inhibition (*32–36*). In addition, JAK-STAT activation has been shown to promote EMT, invasion and metastasis of PCa(*37–41*), further indicating its important role in regulating PCa lineage transition. Collectively, the observed ectopic upregulation of JAK-STAT signaling in the TP53/RB1-deficient and SOX2 overexpression PCa cells raises the intriguing possibility that it may play a crucial role in acquiring lineage plasticity-driven AR therapy resistance.

To dissect the role of JAK-STAT in the enzalutamide resistance associated with TP53/RB1-deficiency, we first knocked-out (KO) TP53 and RB1 in LNCaP/AR cells, a well credentialed enzalutamide-sensitive mCRPC cell model, with CRISPR guides cis-linked with RFP, and generated a stable enzalutamide resistant sgTP53/RB1 clone. Those sgTP53/RB1 cells proliferated significantly quicker when enzalutamide was introduced into the media, in comparison to the sgNT cells expressing GFP (fig.S2A-C). The sgTP53/RB1 cells display clear lineage plasticity as they express significantly increased non-luminal lineage specific genes (fig.S2D), including basal (*KRT5, TP63*), NE-like (*SYP, CHGA, NSE, ASCL1*), and EMT (*CDH2, WNT5A, EPAS1, TGFB, SMAD2*) genes, as well as genes that specify stem cell-like characteristics (*SOX2, NANOG, OCT4*). In the sgTP53/RB1 cells we also observed significant upregulation of many canonical genes activated by the JAK-STAT signaling pathway, which was comparable to the level of JAK-STAT genes induced by SOX2 overexpression (Fig.1B). Consistent with these qPCR results, an increase of H3K27 acetylation (H3K27ac) and H3K4 trimethylation (H3K4me3), as well as a decrease of H3K27 trimethylation (H3K27me3) at the JAK1 gene locus upon depletion of TP53/RB1 was also identified through chromatin immunoprecipitation coupled with qPCR (ChIP-qPCR), indicating a transcriptional upregulation of JAK1 (fig.S3A-C). Interestingly, SOX2 knockout (KO) in the TP53/RB1-deficient cells largely impaired the upregulation of those JAK-STAT signaling genes (Fig.1B), indicating the critical role of SOX2 in the activation of JAK-STAT signaling. This hypothesis is further supported by SOX2 chromatin immunoprecipitation-sequencing (ChIP-Seq) analysis performed on an established enzalutamide resistant mCRPC cell line, CWR-R1, which demonstrated strong and unique SOX2 binding to those canonical JAK-STAT genes in resistant mCRPC, compared to the SOX2 binding in embryonic stem cell line WA01 (fig.S3D, E, raw ChIP-seq data reported in Larischa et al., *Oncogene*, in press).

To determine whether sustained JAK-STAT signaling is required to maintain the resistance in tumor cells with TP53/RB1-deficiency, we KO several of the significantly upregulated JAK-STAT signaling genes in the sgTP53/RB1 cells and observed that depletion of JAK1 and STAT1 blunted the resistant growth of sgTP53/RB1 cells when treated with enzalutamide (Fig.1C, fig.S4A,B). Interestingly, inactivation of those JAK-STAT genes in the cells carrying wildtype TP53 and RB1 did not significantly influence the growth of tumor cells (fig.S4C), suggesting the oncogenic role of JAK-STAT is specific for mCRPC with TP53/RB1-deficiency. These findings were validated *in vivo* in castrated SCID (Severe Combined Immunodeficient) mice treated with enzalutamide, where the depletion of JAK1 and STAT1 largely re-sensitized the sgTP53/RB1 xenografted tumors to enzalutamide treatment (Fig.1D). To dissect the connection between JAK-STAT signaling and lineage plasticity, we examined the expression of canonical lineage marker genes in the sgTP53/RB1/JAK1 cells, which have suppressed JAK-STAT signaling (fig.4D,E), and observed that JAK1 depletion largely attenuated the upregulation of stem-like (*SOX2, NANOG, OCT4, KLF4, NOTCH1*), basal (*KRT5, TP63*), NE-like (*ASCL1, NSE, SYP*) and EMT (*CDH2, TGFB, WNT5A, EPAS1, SNAI1, SMAD2, SMAD3, FN1*) marker genes (Fig.1E-G), which reinforced its crucial role in the acquisition of those non-luminal transcriptional programs. Consistent with an impaired upregulation of EMT as shown by qPCR (Fig.1G), JAK1 depletion also reversed most of the increased migratory and invasive ability of the sgTP53/RB1 cells (fig.S5A-D), supporting the necessity of JAK-STAT signaling in the maintenance of an EMT lineage survival program.

To further explore whether the sustained activation of JAK-STAT signaling is required for the SOX2-promoted lineage plasticity and resistance, we knocked out JAK1 and STAT1 in the LNCaP/AR cells with SOX2-OE and observed that JAK1 and STAT1 depletion almost completely inhibited the resistant growth of LNCaP/AR-SOX2-OE cells when treated with enzalutamide, as shown in both cell proliferation assay (fig.S6A) and CellTiter-Glo assay (fig.S6B). Furthermore, the deactivation of JAK-STAT signaling in the SOX2-OE cells largely attenuated the acquisition of lineage plasticity (fig.S6C), supporting the hypothesis that JAK-STAT activation is required for SOX2-promoted lineage plasticity and AR targeted therapy resistance. Notably, we also observed significantly upregulated expression of stem-like (*SOX2, OCT4, MYC, NOTCH1*), basal (*KRT5*), NE-like (*ASCL1, NSE*) and EMT (*SNAI1, SNAI12, FN1*) marker genes (fig.S6D) in the cells with JAK1 and STAT1 overexpression (JAK1-OE and STAT1-OE), suggesting that JAK-STAT signaling is sufficient to promote the transition to this multilineage and plastic status. The significant upregulation of SOX2 in JAK1-OE cells (fig.S6D, together with Fig1B, fig.S6A-C) also suggested a positive feedback regulation between JAK-STAT signaling and SOX2 in mCRPC tumor cells.

Given the role of JAK-STAT signaling in promoting EMT lineage and AR therapy resistance in our preclinical model, we next examined the impact of JAK-STAT upregulation in various clinical scenarios. We investigated two PCa patient cohorts [The Cancer Genome Atlas (TCGA, Firehose Legacy) and Stand Up To Cancer (SU2C)] and hypothesized that reduced sensitivity to AR targeted therapy would result in relatively higher frequency of copy number amplification and mutations of JAK-STAT genes in the metastatic castration-resistant prostate cancer (mCRPC) compared to hormone-sensitive primary cancers(*42, 43*). Indeed, the frequency of copy number amplification and somatic mutations in canonical JAK-STAT signaling genes were significantly higher in mCRPC (SU2C) compared to hormone naive prostate adenocarcinomas (TCGA) (fig.S7A,B), suggesting a correlation between JAK-STAT upregulation and decreased sensitivity to AR targeted therapy. We next examined both the pathological characteristics and the expression of canonical JAK-STAT genes in the TCGA cohort and discovered that patients with regional lymph nodes metastasis (N1) or high-grade tumors (Gleason score ≥8) have significantly higher JAK-STAT genes expression compared to the patients without regional lymph nodes metastasis (N0) or low-grade tumors (Gleason score≤7) (fig.S7C,D), supporting the role of JAK-STAT in promoting PCa tumorigenesis.

Identification of JAK-STAT signaling as a crucial executor of lineage plasticity-driven resistance raises the hope that appropriate therapeutic approaches targeting this pathway could prevent or overcome AR targeted therapy resistance. For pharmacological inhibition of JAK-STAT signaling, we turned to the specific Jak1 inhibitor, filgotinib. *In vitro* cell viability assays demonstrated that the combination treatment of enzalutamide and filgotinib significantly inhibited the growth of enzalutamide resistant sgTP53/RB1 LNCaP/AR cells (Fig.2A). These results in LNCaP/AR cells were again validated in a second PCa model, the CWR22Pc cells, where JAK1 inhibition by filgotinib significantly inhibited the growth of enzalutamide resistant cells and largely attenuated the upregulation of non-luminal lineage programs (fig.S8A,B). Dose response measurements (IC50) validated that the sgTP53/RB1 cells exhibit less sensitivity to enzalutamide compared to sgNT cells (fig.S8C), while the sgTP53/RB1 cells are much more susceptible to filgotinib treatment compared to sgNT cells (fig.S8D). These *in vitro* results are further supported by *in vivo* xenograft experiments, as the combination treatment of enzalutamide and filgotinib stagnated the growth of enzalutamide resistant sgTP53/RB1 tumors and induced tumor regression compared to either drug alone (Fig.2b). Consistent with the genetic modification results (fig.S6), JAK1 inhibition by filgotinib treatment significantly re-sensitized the SOX2-OE cells to enzalutamide (fig.S8E) and largely attenuated the acquisition of lineage plasticity in these cells (fig.S8F), supporting the hypothesis that JAK-STAT activation is required for the SOX2-promoted lineage plasticity and AR targeted therapy resistance.

**Fig. 2:**
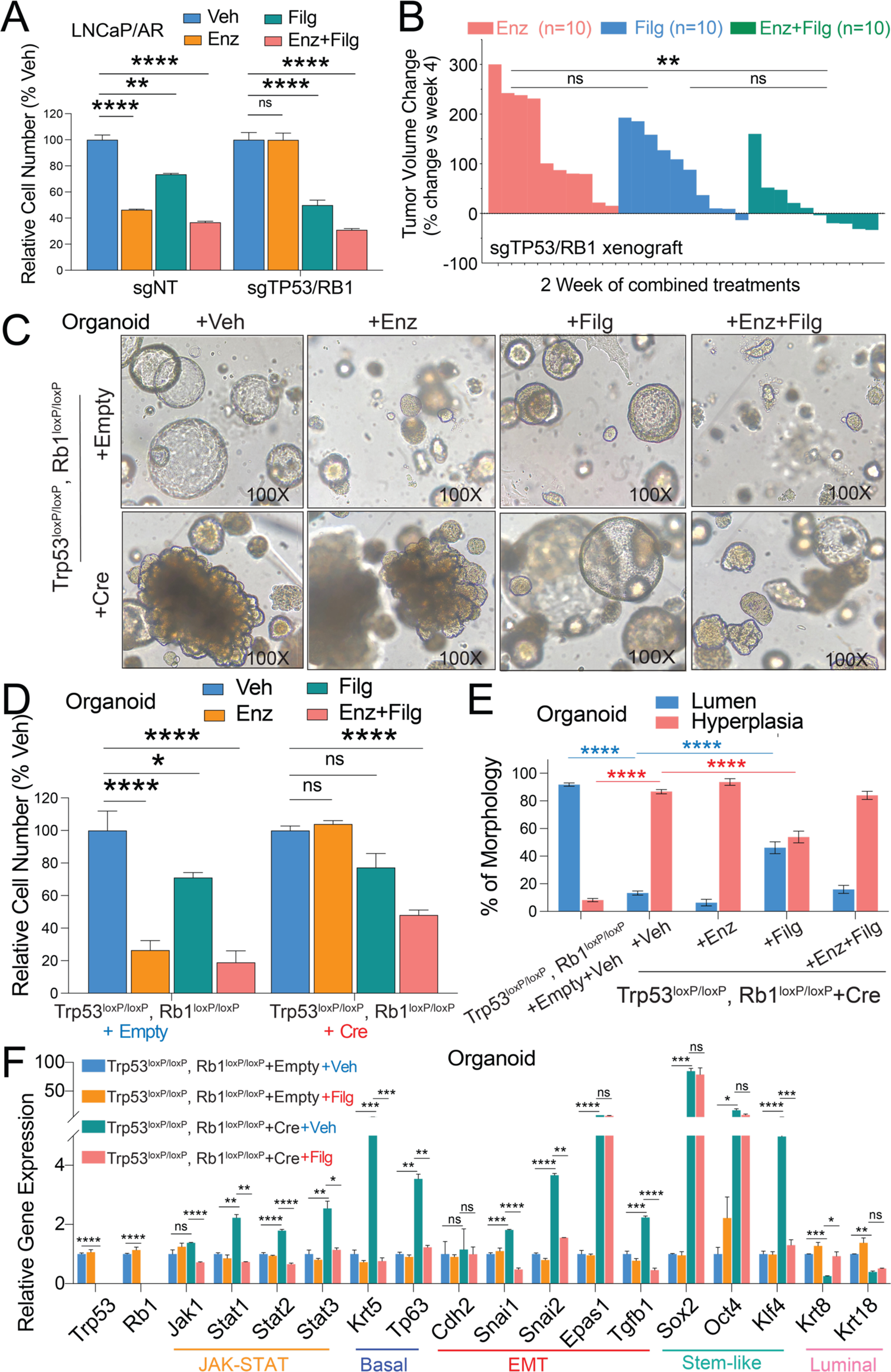
JAK1 inhibitor restores enzalutamide sensitivity in human mCRPC cells and 3D-cultured organoids. (**A**) Relative cell number of LNCaP/AR cells transduced with annotated CRISPR guide RNAs and treated with annotated treatments, normalized to “Veh” group. Enz denotes 10μM enzalutamide, Filg denotes 5μM filgotinib, Enz+Filg denotes the combination of enzalutamide and filgotinib, Veh denotes DMSO treatment with same volume as enzalutamide, for 8 days and cell number were measured by CellTiterGlo assay. (**B**) Waterfall plot displaying changes in tumor size of xenografted LNCaP/AR-sgTP53/RB1 cells after 2 weeks of treatments. All animals were treated with enzalutamide at 10 mg/kg orally 1 day after grafting. Beginning from week 3 of xenografting, animals were randomized into 3 groups and treated with enzalutamide only at 10 mg/kg orally, filgotinib only at 20 mg/kg orally twice daily or the combination of enzalutamide plus filgotinib. mean ± s.e.m. is represented, and p values were calculated using one-way ANOVA. (**C**) Bright field pictures of murine organoids transduced with Cre or empty vector. Organoid were cultured in 3D matrigel and treated with DMSO (Veh), 1μM enzalutamide (Enz), 5μM filgotinib (Filg) or the combination of enzalutamide and filgotinib (Enz+Filg) for 6 days. (**D**) Relative cell number of murine organoids transduced with Cre or empty vectors and treated with annotated treatments for 6 days, normalized to “Veh” group. Treatment’s denotation is same as panel **C**. mean ± s.e.m. is represented, and p values were calculated using two-way ANOVA. (**E**) Percentage of murine organoids display typical lumen or hyperplasia morphology. 3 representative images for each of the lines were counted. Trp53^loxP/loxP^, Rb1^loxP/loxP^-Empty organoids treated with Enz and/or Filg didn’t form typical and large organoid morphology, thus percentage not shown. Treatment’s denotation is same as panel **E**. mean ± s.e.m. is represented, and p values were calculated using two-way ANOVA. (**F**) Relative expression of canonical JAK-STAT and lineage marker genes in 3D-cultured organoids treated with DMSO or Filgotinib. Filg denotes 5μM filgotinib and Veh denotes DMSO treatment with same volume. p values were calculated using two-way ANOVA. For all panels, **** p<0.0001. *** p<0.001. ** p<0.01. * p<0.05. See also **fig.S8**.

To further explore the effect of JAK1 inhibition in a genetically defined model, we utilized our previously generated mouse prostate organoids derived from the Trp53^loxP/loxP^, Rb1^loxP/loxP^ mice after infection with Cre or empty lentivirus (*10*). In contrast to the typical lumen structure, which the Trp53^loxP/loxP^, Rb1^loxP/loxP^+Empty organoids formed in 3D culture, the Trp53^loxP/loxP^, Rb1^loxP/loxP^+Cre organoids displayed a hyperplastic morphology, where the organoid cells formed a solid ball and finger-type structures invading the surrounding matrigel (Fig.2C), indicating an invasive phenotype. These Trp53^loxP/loxP^, Rb1^loxP/loxP^+Cre organoids were significantly resistant to enzalutamide treatment compared to the Trp53^loxP/loxP^, Rb1^loxP/loxP^+Empty controls (Fig.2C,D), but responded well to the combination treatment of enzalutamide and filgotinib (Fig.2C,D). Remarkably, we also observed a significant number of the Trp53^loxP/loxP^, Rb1^loxP/loxP^+Cre organoids re-established a classic lumen-like structure when treated with filgotinib compared to vehicle treated group (Fig.2C,E), indicating that JAK1 inhibition by filgotinib impairs the upregulation of non-luminal transcriptional programs due to Trp53 and Rb1 depletion. This hypothesis is further supported by qPCR results showing attenuated upregulation of the basal, EMT and stem cell-like lineage genes (Fig.2F) in those organoids.

Since JAK1 depletion largely re-sensitized the sgTP53/RB1 cells to enzalutamide treatment, we next sought to determine whether JAK-STAT signaling is specifically required for the therapy resistance of heterogeneous subclones with lineage plasticity. Considering that the analysis of bulk cell RNA-Seq represents an average of gene expression across a large population of potentially heterogeneous cells expressing various lineage transcriptional programs, we performed single cell RNA-Seq and transcriptomic analysis using the LNCaP/AR sgNT, sgTP53/RB1, and sgTP53/RB1/JAK1 cell lines treated with enzalutamide or vehicle for 5 days (∼10,000 cells per group). As expected, clustering of the sequenced cells was primarily driven by the genetic modifications and treatments of these cells (Fig.3A-C). Interestingly, the majority of both the sgNT and sgTP53/RB1/JAK1 cells are clearly separated by the different treatments (enzalutamide vs vehicle, as shown in Fig.3A, C), while the sgTP53/RB1 cells do not display a similar separation (mixed population shown in Fig.3B), indicating that majority of the sgTP53/RB1 cells exhibit enzalutamide resistance. Since AR antagonists can prohibit PCa cell growth by promoting cell cycle arrest (*10, 44*), we performed cell cycle distribution prediction analysis of each single cell using the scRNA-Seq data and observed a dramatically increased cell cycle arrest occurring in the sgNT cells treated with enzalutamide, as nearly 80% of the cells were in the G1 phase compared to less than 30% of the cells in the vehicle treated group (Fig.3A,D). In contrast, enzalutamide treatment does not increase the population of cells in G1 phase in the sgTP53/RB1 cell group, supporting that majority of the sgTP53/RB1 cells are resistant to enzalutamide-caused cell cycle arrest (Fig.3B,D). Remarkably, JAK1 depletion in the sgTP53/RB1 cells significantly increased the percentage of cells entering G1 phase upon the treatment of enzalutamide compared to the vehicle treated group (Fig.3C,D), suggesting that JAK1 depletion re-sensitized those sgTP53/RB1 subpopulations to enzalutamide treatment. Notably, JAK1 depletion didn’t impair the proliferation of sgTP53/RB1 cells when treated with vehicle (Fig3C,D), suggesting JAK-STAT’s specific role in mediating AR targeted therapy resistance.

**Fig. 3.**
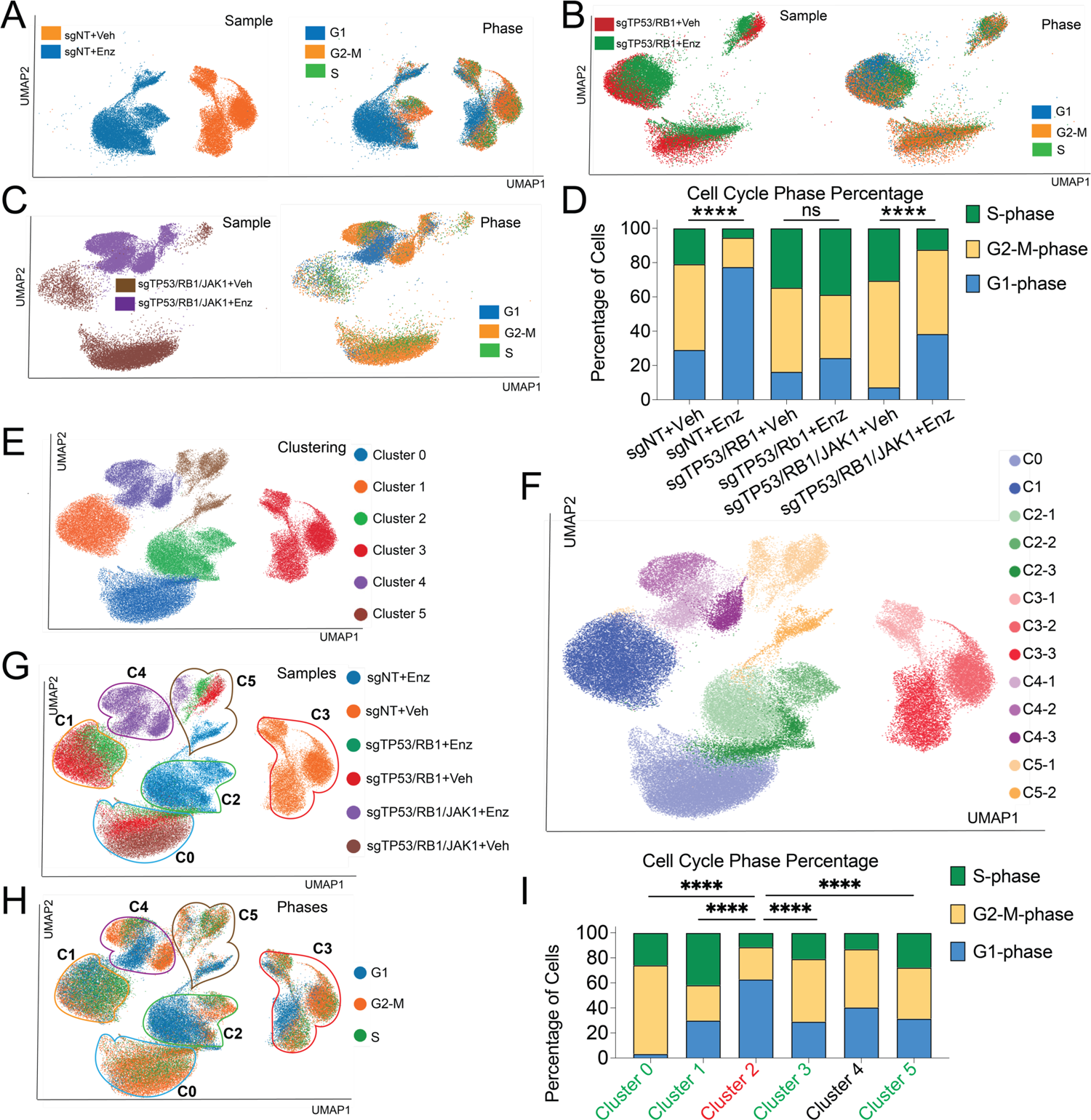
JAK-STAT is required for the survival of resistant subclones of cells in TP53/RB1-deficient mCRPC. (**A-C**) UMAP plots of single cell transcriptomic profiles of LNCaP/AR cells transduced by annotated CRISPR guide RNAs, treated with vehicle (DMSO) or 10µM enzalutamide for 5 days. a, LNCaP/AR-sgNT (Veh n=14268, Enz n=15149), (**B**) LNCaP/AR-sgTP53/RB1(Veh n=12267, Enz n=9850), and (**C**) LNCaP/AR-sgTP53/RB1/JAK1(Veh n=25200, Enz n=11096). The left panels were colored according to sample origin while cells in the right panels were colored by predicted cell cycle phase. (**D)** Bar plot presents the percentage distribution of each single cells in different cell cycle phases in each sample. p-values are calculated with Fisher’s Exact Test. *** p<0.001. (**E**) Single-cell profile of LNCaP/AR cells based on clustering. UMAP plot of single cells colored by unsupervised clustering of 6 subsets is presented. (**F**) Single-cell profile of LNCaP/AR cells based on sub-clustering. UMAP plot of single cells colored by unsupervised clustering of 13 sub-clusters is presented. (**G**) Single-cell profile of LNCaP/AR cells transduced with annotated CRISPR guide RNAs and treated with vehicle or enzalutamide. UMAP plot of single cells colored by samples is represented. Area and number of clusters in panel **E** is highlighted with color circles. (**H**) Single-cell profile of LNCaP/AR cells based on cell cycle states. UMAP plot of single cells colored by cell cycle prediction is presented. Area and number of clusters in panel **E** is highlighted with color circles. (**I**) Bar plot presents the percentage distribution of each single cells in different cell cycle phases in each of the 6 clusters. p-values are calculated with Fisher’s Exact Test. For all panels, **** p<0.0001,*** p<0.001,** p<0.01, * p<0.05. ns: not significant. See also **fig.S9** and **table S2**.

To further assess the dynamics of the resistance in TP53/RB1-deficient mCRPC at single cell resolution, we next investigated whether AR signaling was fully or partially restored in the resistant subclones of cells, as previously shown in many subtypes of resistant mCRPC(*4*). Not surprisingly, sgNT+Veh group consisted of the greatest number of cells expressing canonical AR target genes (the well-established AR-Score genes, table S2) and inhibition of their expression was subsequently verified upon enzalutamide exposure (fig.S9). In contrast, both the sgTP53/RB1-Veh and sgTP53/RB1-Enz groups predominantly lack the expression of the AR target genes, further supporting the dominant role of AR-independent transcriptional survival programs in those resistant cells (fig.S9). Interestingly, the expression of canonical AR target genes was largely re-established in many cells belonging to the sgTP53/RB1/JAK1+Veh group (two thirds of the AR-Score genes, table S2), compared to the sgTP53/RB1+Veh group (fig.S9A), suggesting a partial restoration of AR signaling and AR dependency, as well as an elevated cellular heterogeneity, among the sgTP53/RB1/JAK1 cells.

Since TP53/RB1 deficiency, as well as the deactivation of JAK-STAT signaling, significantly altered the expression of lineage-specific transcriptional programs in the bulk cell population (Fig.1E-G), we surveyed how JAK-STAT signaling affected the acquisition of lineage plasticity at single cell resolution. To characterize the lineages of different cell populations, we performed unsupervised graph clustering (Uniform Manifold Approximation and Projection, UMAP)(*45*) and identified 6 distinct cell subsets labeled as Cluster 0-5, with further partitioning to 13 sub-clusters (Fig.3E-F). Consistent with the significant transcriptomic changes caused by TP53/RB1/JAK1 modification, 5 of the 6 clusters (clusters 0-4) predominantly overlapped with the clusters identified by genetic modifications and treatment groups (Fig.3G). Intriguingly, Cluster 5 is a mixture of a small fraction of cells from five groups: sgNT+Enz, sgTP53/RB1+Veh, sgTP53/RB1+Enz, sgTP53/RB1/JAK1+Veh, and sgTP53/RB1/JAK1+Enz (Fig.3E-G). To examine the cell proliferation status of these clusters, we overlapped the transcriptomic-based clustering with cell cycle prediction (Fig.3H). Interestingly, cells within the Cluster 0, 1, 3, and 5 remain proliferative (termed the “winner” clusters, Fig.3I), whereas Cluster 2 contains a much higher percentage of cells in cell cycle arrest (termed the “loser” cluster, Fig.3I). Lastly, the cells within Cluster 4 express elevated levels of cell cycle phase heterogeneity, a finding that will be expounded upon later (Fig.3H).

To explore the lineage characterization of these clusters, we sought to determine which of these clusters culminated in the expression of various lineage-specific transcriptional programs. We probed the well-established AR-Score gene signature and five lineage-specific gene signatures(*10, 11, 46–48*) (table S2) and analyzed the expression of the genes (z-score) comprising these signatures across all six clusters as well as samples of single cell subsets (Fig.4A-C). In congruence with the luminal epithelial cell lineage of the original LNCaP/AR cells, Cluster 2 and Cluster 3, consisting predominantly of cells originating from the sgNT groups, represent the two clusters expressing the highest level of the luminal gene expression (Fig.4A-D). Since the survival of these luminal epithelial cells depends on AR signaling, most of Cluster 2 cells, while retaining their luminal lineage, displayed loss of AR signaling gene expression and entered cell cycle arrest upon enzalutamide treatment (Fig4A-E). Notably, the most substantial proportion of Cluster 0 and 1, consisting primarily of cells originating from the sgTP53/RB1 groups, expressed the lowest level of luminal gene signature and relatively high levels of non-luminal lineage gene signatures compared to Cluster 3 (predominately sgNT+Veh), including the EMT, stem-like, basal, and NE-like lineage gene signatures (Fig.4A-I). Interestingly, Cluster 0 and 1 also contained a substantial proportion of cells from the sgTP53/RB1/JAK1+Veh group which maintained the expression of non-luminal transcriptional programs (Fig.4B-I), supporting the hypothesis that the deactivation of JAK-STAT signaling does not impair the general survival of those subclones in the absence of AR targeted therapy (enzalutamide) (Fig.4B-C, Fig.3I). However, enzalutamide treatment dramatically diminished the survival of sgTP53/RB1/JAK1 subclones and the expression of stem-like, EMT and basal multilineage programs, suggesting that JAK-STAT inactivation restored AR-dependency and impaired the lineage plasticity and AR therapy resistance of those subclones (Fig.4C). This hypothesis is further supported by the partially restored AR signaling in those sgTP53/RB1/JAK1 subclones (fig.S9). Interestingly, although JAK-STAT has been shown to be required for the resistance of lineage plastic subclones expressing multilineage programs, including stem-like, EMT, and basal lineages (Fig.4C), the deletion of JAK1 did not significantly impair the resistance of subclones expressing an NE-like lineage program (Fig.4C), indicating that JAK-STAT is specifically required for the de-differentiation to a stem-like and lineage plastic status, rather than following re-differentiation to the NE-like lineage in some subclones.

**Fig. 4:**
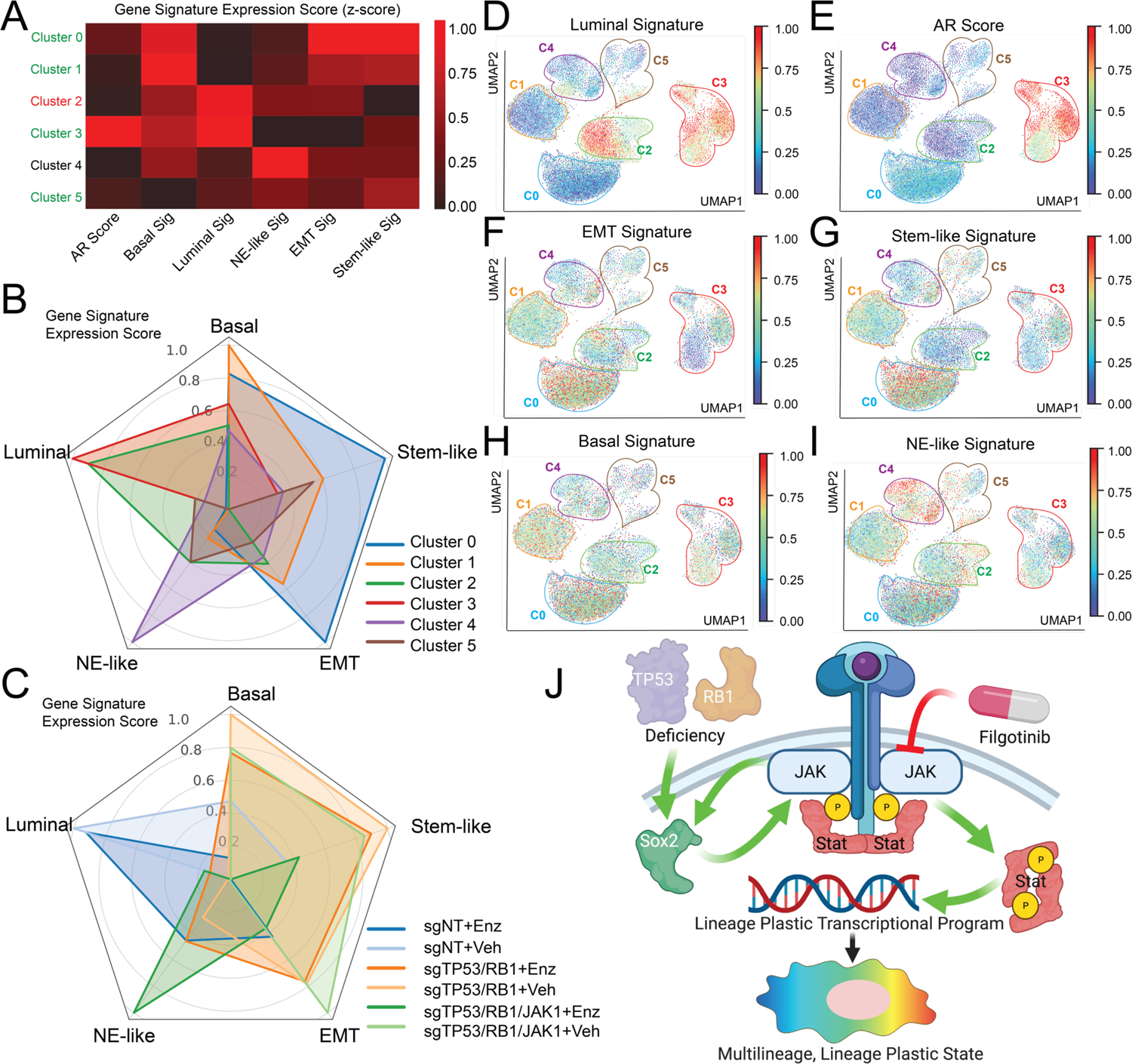
JAK-STAT signaling is required for the maintenance of lineage plastic subsets of mCRPC cells. (**A**) Heatmap represents the lineage scores of canonical lineage marker gene signatures in cell clusters. Winner clusters (without increased cell cycle arrest) is highlighted in green and loser clusters (with increased cell cycle arrest) is highlighted in red. (**B**) Radar plot represents the lineage scores and distribution of different cell clusters. (**C**) Radar plot represents the lineage scores and distribution of different samples. For panel **A-C,** lineage scores were scaled to 0-1 across all clusters. (**D**) UMAP plot of single cell transcriptomic profiles colored by luminal gene signature score (z-score) for each cell (dot). (**E**) UMAP plot of single cell transcriptomic profiles colored by AR gene signature score (z-score) for each cell (dot) of LNCaP/AR cells transduced with annotated CRISPR guide RNAs and treated with vehicle or enzalutamide. (**F**) UMAP plot of single cell transcriptomic profiles colored by EMT gene signature score (z-score) for each cell (dot). (**G**) UMAP plot of single cell transcriptomic profiles colored by Stem cell-like gene signature score (z-score) for each cell (dot). (**H**) UMAP plot of single cell transcriptomic profiles colored by basal gene signature score (z-score) for each cell (dot). (**I**) UMAP plot of single cell transcriptomic profiles colored by NE-like gene signature score (z-score) for each cell (dot). For panel **A-F,** distribution area of each LNCaP/AR cell line sample numbers are labeled with black and each of the Clusters are labeled in color circles. Color density of each cell is scaled by the color bar. For all panels, lineage scores were scaled to 0-1 across all cells. (**J**) Schematic describing that JAK-STAT transcriptionally upregulated in the mCRPC cells with TP53/RB1 deficiency and ectopic SOX2 activity, created with BioRender.com. See also **fig. S9-10** and **table S2**.

As the data derived from our single cell sequencing revealed clear heterogeneity within the enzalutamide treated sgTP53/RB1/JAK1 cells (Fig.3E-F), we continued to explore the lineage characterization of the subclusters of Cluster 4 (Fig.3F), which predominantly contains cells originating from the sgTP53/RB1/JAK1+Enz group (Fig.3E,G). Interestingly, the three subclusters of Cluster 4 expressed diverse levels of the JAK-STAT genes (fig.S10), presumably because the JAK-STAT signaling was not fully deactivated in a proportion of JAK1-KO cells due to compensatory signaling from other JAKs and STATs. The JAK-STAT signaling heterogeneity detected in Cluster 4 provided a unique opportunity to decipher whether the successful deactivation of JAK-STAT signaling in a subpopulation of cells would impair lineage plasticity and therapy resistance. Compared to the other subclusters of Cluster 4, Cluster 4-3, contained the “outlier” cells, which originate from the sgTP53/RB1/JAK1 group yet partially maintain JAK-STAT signaling (Fig.3F, fig.S10B-I), likely due to a compensatory activation of JAK-STAT signaling. Remarkably, Cluster 4-3 cells maintained the expression of multilineage transcriptional programs, including stem-like, EMT, basal and NE-like lineages, and remained proliferative even in the presence of enzalutamide (Fig.3F, H, Fig.4F-I, fig.S10A). By contrast, Cluster 4-1 contains cells expressing decreased levels of the multilineage transcriptional programs (Fig3F, Fig4F-I) and were highly responsive to the treatments of enzalutamide (Fig.3F, H, fig.S10A), suggesting an impaired AR therapy resistance in those subclones. Interestingly, Cluster 4-2 is a subclone which only maintains the NE-like genes expression, rather than the lineage plastic and multilineage transcriptional programs (Fig.3F, Fig.4I). Similarly, Cluster 4-2 also maintained active cell proliferation despite the treatment of enzalutamide (Fig.3F, H, fig.S10A), further supporting the hypothesis that JAK-STAT signaling is not required for the resistance of subclones that have already re-differentiated to an NE-like lineage. The juxtaposition between Cluster 4-1 (fully deactivated JAK-STAT signaling with impaired lineage plasticity), Cluster 4-3 (partially maintained JAK-STAT signaling and multilineage programs) and Cluster 4-2 (fully deactivated JAK-STAT signaling with impaired lineage plasticity but maintained NE-like lineage) further supports the crucial role of JAK-STAT signaling in maintaining the AR therapy resistance of stem-like and lineage plastic subclones expressing multilineage transcriptional programs, rather than the subclones fully re-differentiated to an NE-like lineage. Collectively, these results suggest that JAK-STAT signaling is a crucial executor of lineage plasticity-driven AR targeted therapy resistance in the TP53/RB1-deficient mCRPC (Fig.4J).

Emerging evidence demonstrates that lineage plasticity represents an important mechanism for conferring targeted therapy resistance in various cancers, particularly prominent in cancers where the molecular target of therapies are the lineage-specific survival factors, including ER-positive breast cancer, EGFR-mutant lung cancer, BRAF-mutant melanoma, and AR-dependent PCa (*7, 9–16, 19–22*). In the case of PCa, however, it is not fully understood whether the differentiated luminal tumor cells acquire lineage plasticity-driven resistance through reverting back (de-differentiating) to a multi-lineage, stem cell-like state and then re-differentiating to alternative lineages, or through direct trans-differentiation to a distinctively new lineage (*10, 18, 49*). Another intriguing feature of lineage plasticity-driven targeted therapy resistance is the elevated levels of intratumoral heterogeneity(*50*). However, average gene expression signals analyzed from bulk cell populations in previous studies may have masked the intratumoral heterogeneity, which could obstruct the dissection of the molecular mediators required either for lineage plasticity or for a specific lineage program such as NE-like lineage. Thus, the identification of heterogeneous TP53/RB1-deficient tumor cell subpopulations expressing various lineage programs through single cell transcriptomic analyses illuminates these once hidden details and represents a major insight into this work. Here, by using single cell transcriptomic profiling, we showed that a vast majority of the TP53/RB1-deficient tumor cells acquire lineage plasticity by transitioning to a lineage plastic, multi-lineage, stem cell-like, and AR independent state with concurrent expression of an EMT transcriptional program *in vitro.* Importantly, our data also suggested that ectopic JAK-STAT activation is required for the AR therapy resistance of those de-differentiated, stem-like cells expressing lineage plastic and multilineage transcriptional programs, rather than the cells having undergone complete NE-like trans-differentiation.

Various genetic and transcriptional aberrations have been connected to the acquisition of lineage plasticity in PCa, including, but not limited to, the aberrations of PTEN, BRN2, FOXA1, N-Myc, PEG10, CHD1, REST, and BRG1 (*7, 11–16*). Interestingly, many of those cases involve the “hijacking” of stem-like, pluripotency, or epigenetic regulation programs, such as SOX2, SOX11, EZH2, and the SWI/SNF complex (*9, 10, 13–15, 18*). Although the role of JAK-STAT signaling pathway in regulating cell fate decision, stem cell self-renewal, and multilineage differentiation has been well documented (*23–25*), its potential function in mediating lineage plasticity-driven AR therapy resistance remained largely unclear. Here, our results indicate a significant role of JAK-STAT signaling in TP53/RB1 deficient tumor cells, whereby the activation of the pathway promotes the transition of luminal epithelial cells to a multilineage, stem cell-like, and EMT status. These results are consistent with previous findings that JAK-STAT promotes EMT transition and tumor metastasis, often through the induction of pluripotency signaling transduction, in uveal melanoma, colorectal, breast, head and neck, and prostate cancers(*37, 38, 51–54*).

Although the oncogenic roles of JAK-STAT signaling, including the activation of STAT1, STAT3 and STAT5, have been widely corroborated in various cancers including PCa, the exact biological consequence of constitutive activation of those STAT proteins in tumorigenesis is highly context specific(*28*). For example, IL6-induced STAT3 activation has been reported to promote PCa NE-like differentiation and cell cycle arrest(*29–31*), while also protecting PCa cell from apoptosis caused by AR inhibition(*32–36*). Similarly, despite its known function in mediating antitumor immune surveillance, ectopic expression of STAT1 confers radioresistance in squamous cell carcinoma and chemotherapy resistance in colon cancer(*28, 55, 56*). Here, we showed that the JAK-STAT signaling activation, partially in a STAT1-depedent manner, is required for the lineage plasticity-driven AR therapy resistance in TP53/RB1-deficient tumors, but not for the tumor cells which have completely re-differentiated to an NE-like lineage. Our data also indicated that the JAK-STAT signaling enables the expression of EMT transition lineage program which promotes a metastatic phenotype of the resistant cells. This mechanism parallels the behavior of p53-deficient esophageal tumors, which demonstrates that ectopic upregulation of STAT1 promotes tumor metastasis and invasion(*57*). Such correlations between STAT1 and the activation of EMT transcriptional programs have also been observed in triple negative breast cancer and colon cancer (*55, 58*). Furthermore, our single cell analysis showed that the JAK-STAT signaling was not completely deactivated in a subset of cells originating in the sgTP53/RB1/JAK1 group, which suggests the possibility that various JAK and STAT proteins function in a collaborative and compensatory network.

It is important to place our model of how JAK-STAT signaling is hijacked to promote lineage plasticity, EMT transition, and resistance in the context of TP53 and RB1 deficiency (*42*). Accumulating evidence suggests a connection between JAK-STAT activation and TP53/RB1 alterations in various cancers. In EGFR-mutant lung cancer, concurrent TP53/RB1 alterations define a subset of tumors with small cell lung cancer (SCLC) transformation, which contains significantly enriched mutation frequencies in the JAK-STAT signaling pathway(*59*). However, others have documented an inverse correlation between wildtype TP53 and JAK-STAT activation. For example, wildtype TP53 is reported to inhibit the transcriptional activity of both STAT3 and STAT5 in myeloproliferative neoplasms (MPNs)(*60, 61*), the latter of which prevents STAT5 from binding to lineage specific factors which drive differentiation (*62*). These results are consistent with our finding that the inactivation of JAK-STAT signaling dramatically impairs the proliferation of resistant cells with TP53/RB1-deficiency, while not affecting the cells with intact TP53 and RB1 (fig.S4C). Therefore, it is critical to consider the genomic status of TP53/RB1 when correlating JAK-STAT activation with the clinical outcome of AR therapy responses, as the JAK-STAT activation in patients with wildtype TP53/RB1 may not be a consequence of lineage “hijacking,” but rather cytokine-induced immune response. However, this extrapolation would require the analysis of a much larger cohort of mCRPC patients carrying TP53/RB1 alterations.

Despite the clinical success of AR targeted therapies in controlling mCRPC, acquired resistance to these treatments universally develops and largely impairs the clinical outcome of patients with mCRPC. Although lineage plasticity-driven resistance has been suggested as a substantial mechanism conferring resistance and several underlying mechanisms have been revealed, effective therapeutic approaches for patients with resistant mCRPC driven by lineage plasticity are still not available. Although our previous discovery reveals an important role of SOX2 in mediating lineage plasticity(*10*), direct pharmacological inhibition of SOX2 is not currently feasible, underscoring the unmet need to develop novel combination therapies targeting lineage plasticity in this subtype of lethal mCRPC with TP53/RB1-deficiency. Here, using various human mCRPC cell models and 3D-cultured organoid model, we demonstrated the efficacy of JAK1 inhibitor filgotinib, in combination with enzalutamide, to overcome the lineage plasticity-driven resistance in TP53/RB1-deficient mCRPC. These results may provide strong rationale for future clinical trials designed to target JAK-STAT signaling for overcoming lineage plasticity-driven AR targeted therapy resistance.

## Supporting information

Gene Set Enrichment Analysis (GSEA) results show significantly changed signaling pathways

AR Score and lineage specific signatures gene lists

## Acknowledgments

We thank the SU2C, TCGA, cBioPortal.org, and Firehose Legacy (Firebrowse.org) for providing genomic and transcriptomic data. We thank Dr. Charles L. Sawyers for some of the reagents used in this study. We thank Drs. Kathryn O’Donnell, Michael Buszczak and Ganesh Raj for helpful discussion.

## Author contributions

S.D. and P.M. conceived the project.

S.D., CS.W, YG.W. and P.M. designed, conducted experiments and interpreted data.

S.D. and P.M. co-wrote the manuscript.

L.M. and N.J. edited the manuscript.

S.D., CS.W. and XL.L. conducted all genetic and pharmaceutical inactivation of JAK-STAT signaling in all the *in vitro* assays.

S.D., CS.W. and N.J. performed all in vivo xenograft experiments.

XL.L., K.R., J.J. and P.M. performed all organoids experiments.

YR.X., J.G. and S.D. performed all ChIP and migration assay experiments.

S.D., CS.W., XL.L., C.R.T. and V.A. performed all qPCR and western blot experiments.

LF.X., UG.L., JT. H. and P.M. performed clinical data analysis.

B. L. and JF. Y. conducted the library preparation and sequencing of single cell RNA-Seq.

YG.W. and C.A. performed bioinformatic analysis for bulk RNA-Seq and YG.W. performed analysis for single RNA-Seq.

T.W. and P.M. oversaw the bioinformatic analysis.

D.J.V. performed the SOX2-ChIP-Seq and analysis.

ZQ. X. conducted the deposit of bioinformatic data.

P.M. are the corresponding authors of this manuscript.

## Data and materials availability

Further information and requests for resources and reagents should be directed to and will be fulfilled by the Lead Contact, Dr. Ping Mu. All cell lines, plasmids and other reagents generated in this study are available from the Lead Contact with a completed Materials Transfer Agreement if there is potential for commercial application. All the described bulk RNA-seq data and single cell RNA-seq data have been deposited in the Gene Expression Omnibus under the accession numbers GSE175975.

## Supplementary Materials

### Materials and Methods

#### Human cell line and mouse organoid culture

LNCaP/AR and CWR22Pc prostate cancer cell lines were generated and maintained as previously described(*7, 10, 63, 64*). LNCaP/AR and CWR22Pc cells were cultured in RPMI 1640 medium supplemented with 10% fetal bovine serum (FBS), 1% L-glutamine, 1% penicillin-streptomycin, 1% HEPES, and 1% sodium pyruvate (denoted as normal culture medium). LNCaP/AR cells were passaged every 3-5 days at a 1:6 ratio, CWR22Pc cells were passaged every 3-5 days at 1:3 ratio. When treated with 10 µM enzalutamide and/or 5 µM filgotinib, LNCaP/AR cells were cultured in RPMI 1640 medium supplemented with 10% charcoal-stripped serum (denoted as CSS medium). All cell cultures were assessed for mycoplasma monthly via the highly sensitive MycoAlertTM PLUS Mycoplasma Detection kit from Lonza (Cat #LT07-710). Cell line identification was validated each year through the human STR profiling cell authentication provided by the UT Southwestern genomic sequencing core and compared to ATCC cell line profiles. Trp53^loxP/loxP^, Rb1^loxP/loxP^ murine organoids were generated from Trp53^loxP/loxP^, Rb1^loxP/loxP^ mice as previously described(*10*). The organoids are cultured in 3D Matrigel according to established protocol (*65, 66*). The organoids are split at 1:3 ratio every 6 days by trypsin or sterile glass pipette. Organoids were transduced with lentivirus constructs of either Cre or DsRed (empty) as control and selected with 1 μg/ml puromycin for 5 days, 2 days post transduction as previously described(*10*). When treated with 1μM enzalutamide and/or 5μM filgotinib, these organoids were cultured in typical murine organoid medium supplemented with drugs (*65, 66*)

#### CRISPR model generation

Lentiviral transduction of cells for guide RNA experiments was performed as previously described with some modifications(*7, 10, 67*). Lentiviral virus was used for CRISPR-based knockout of TP53, RB1, and all the other genes modified in the manuscript. CRISPR-mediated gene modification was performed as previously described(*7*). Specifically, LNCaP/AR cells were seeded at 400,000 cells per well in 2 ml of media in 6-well plates. The next day, media was replaced with media containing 50% of virus and 50% of fresh culture medium, along with 5 μg/ml polybrene. The lentiviral virus containing media was removed after 24 hours and replaced with normal culture medium. Three days post transduction, the cells were selected with 2 μg/ml puromycin for 4 days or 5 μg/ml blasticidin for 5 days. For cells with double colors, transduced cells were further sorted by Flow Cytometer for double positive population. All shRNAs and related constructs have been previously described(*7, 10, 67*). Human DYKDDDDK (Flag)-tagged-SOX2 expression lentivirus (cat #337402) was purchased from Qiagen and used for direct cell transduction, following the manufacturer’s instruction. The All-In-One lentiCRISPR v2 (Addgene Plasmid #52961), LentiCRISPRv2GFP (Addgene Plasmid #82416), LentiCRISPRv2-mCherry (Addgene Plasmid #99154), pLKO5.sgRNA.EFS.RFP (Addgene Plasmid #57823), pLKO5.sgRNA.EFS.GFP, lentiCas9-Blast (Addgene Plasmid #52962) plasmids were used to generate the CRISPR and guide RNAs targeting TP53, RB1, JAK1, and all the other genes modified in the manuscript. The guide RNA constructs with empty space holder served as the sgNT control. The guide RNAs were designed using the online CRISPR designing tool at Benchling (https://benchling.com). The sequences of sgRNAs are listed below:

sgRB1-F: CACCGATAGGCTAGCCGATACACTG

sgRB1-R: AAACCAGTGTATCGGCTAGCCTATC

sgTP53-F: CACCGCCATTGTTCAATATCGTCCG

sgTP53-R: AAACCGGACGATATTGAACAATGGC

sgJAK1-F: CACCGATCTTCTATCTGTCGGACA

sgJAK1-R: AAACTGTCCGACAGATAGAAGATC

sgSTAT1-F: CACCGTTATGATGACAGTTTTCCCA

sgSTAT1-R: AAACTGGGAAAACTGTCATCATAAC

sgSTAT2-F: CACCGGTGCAGCTGATCCTGAAAG

sgSTAT2-R: AAACCTTTCAGGATCAGCTGCACC

sgSTAT3-F: CACCGACAGCTTCCCAATGGAGCTG

sgSTAT3-R: AAACCAGCTCCATTGGGAAGCTGTC

#### *in vivo* xenografts experiment

All animal experiments were performed in compliance with the guidelines of the Animal Resource Center of UT Southwestern, similarly as previously described (*7*). LNCaP/AR *in vivo* xenograft experiments were conducted by subcutaneous injection of 2 × 10^6^ cells (100 μl in 50% Matrigel and 50% growth media) into the flanks of castrated male SCID mice on both sides. For experiment in Fig.1D, daily gavage treatment with 10 mg/kg enzalutamide or vehicle (1% carboxymethyl cellulose, 0.1% Tween 80, 5% DMSO) was initiated one day after the injection. Once tumors were noticeable, tumor size was measured weekly by digital caliper. For experiments in Fig 2B, 10 mg/kg enzalutamide (daily) and/or 20 mg/kg filgotinib (twice daily) were given after 3 weeks of enzalutamide alone administration, when tumors averaged around 200 mm^3^ in size. Enzalutamide was purchased from the Organic Synthesis Core Facility at MSKCC. Filgotinib is commercially available from MedChem Express.

#### Cell Dose Response Curve, Growth, Viability, and FACS-based Competition Assays

Cell growth assay, viability assay, dose repones curve and competition assay were conducted as previously described (*7*). Specifically, for viability assay and dose response curve, 4000 LNCaP/AR cells were seeded in 96-well plate and treated with different dosages of treatments for 8 days before performing the assay, then cell viability were measured by CellTiter-Glo luminescent cell viability assay (Promega cat #7570) according to manufacture protocol. For cell growth assay, LNCaP/AR (10,000 cells per well) or CWR22Pc (50,000 cells per well) cells were seeded in a 24-well cell culture plate, in FBS medium (CWR22Pc) or CSS medium (LNCaP/AR) and treated with enzalutamide (10 μM for LNCaP/AR, 1 μM for CWR22Pc) or vehicle (DMSO) for 7 days (LNCaP/AR) or 4 days (CWR22Pc) and cell numbers were counted. Cell growth assays were conducted in triplicate and mean ± S.E.M. were reported. For organoid growth assay, 2000 murine organoid cells were seeded in 3D Matrigel (per 50 µl sphere) in murine organoid media (*65, 66*)with enzalutamide and/or filgotinib for 6 days. Matrigel was washed away with cell recovery medium (Corning, cat #354253) and organoids were separated into single cell suspension by trypsin, then cell numbers were counted, and the relative cell growth (treatments/veh) was calculated. For FACS-based competition assay, the competition cell mixture of ∼20% sgTP53/RB1-RFP cells and ∼80% sgNT-GFP cells was treated with 10 μM enzalutamide and the percentage of RFP positive cells were measured by FACS on day 0, day 4, day 8. Relative cell number fold change was calculated and normalized to veh treated group as previously described (*7*).

#### Boyden chamber migration and invasion assays

20,000 LNCaP/AR cells were resuspended in serum free RPMI, seeded in the upper transwell insert (Corning cat #353097). RPMI with 10% serum was added to the lower chamber as a chemoattractant. After 60 h incubation, cells that migrated to the lower side of the transwell insert were fixed with PFA, stained with 1% crystal violet. Images were acquired on Leica DMi8 inverted microscope. 9 representative images of each group were used to quantify the migrated cell numbers using ImageJ. For invasion assay, the inserts were coated with a layer of extracellular matrix (ECM) gel, Matrigel (Corning, Cat# 354234), before plating. The stock Matrigel (10 mg/ml) was thawed overnight at 4 °C and then diluted in cold serum-free RPMI to a working amount of 30 μg per insert. Each insert was coated with 100 μl of diluted Matrigel and incubated 1 h at 37 °C in a humidified atmosphere in the presence of 5% CO_2_. Following incubation of the gel layer, cells were plated at the same density and in the same manner as described in the migration section. After allowing 60 h for invasion, cells were fixed, stained with 1 % crystal violet, and quantified using the same method as migration assay as described above.

#### Gene expression detection by qPCR, Western Blot

qPCR and western blot experiments were conducted as previously described (*7, 10, 67*). Specifically, Total RNA from cells was extracted using Trizol (Ambion, Cat 15596018) following manufacturer’s instructions. cDNA was made using the SuperScript™ IV VILO™ Master Mix with ezDNase™ Enzyme (Thermo Fisher, 11766500) following manufacturer’s instructions, with 200 ng/µl RNA template. 2X PowerUp™ SYBR™ Green Master Mix (Thermo Fisher, A25778) was used in the amplification of the cDNA. Assays were performed in triplicate and normalized to endogenous β-Actin expression. For western blot, proteins were extracted from whole cell lysate using RIPA buffer, then measured with Pierce BCA Protein Assay Kit (cat #23225). Protein lyses were boiled at 95°C for 5 minutes and run on the NuPAGE 4-12% Bis-Tris gels (Invitrogen, Cat #NP0323). Transfer was conducted at 4°C for 1 hour at 100 volts. Membranes were blocked in 5% non-fat milk for 15 minutes prior to addition of primary antibody and washed with 1X TBST (10X stock from Teknova, T9511). Antibodies for western blot are listed: JAK1 (Cell Signaling Technology, Cat # 3332S), STAT1 (Cell Signaling Technology, Cat #9172S), STAT2 (Cell Signaling Technology, Cat # 93130T), STAT3 (Cell Signaling Technology, Cat # 9139T), p-STAT1 (Cell Signaling Technology, Cat # 9167S), Rb (Cell Signaling Technology, Cat # #5230), P53(Leica Biosystems, Cat# NCL-p53-DO1), Actin (Cell Signaling Technology, cat #4970).

Human qPCR primers for gene expression detection are listed:

JAK1 (F-GAGACAGGTCTCCCACAAACAC; R-GTGGTAAGGACATCGCTTTTCCG)

JAK2 (F-CCAGATGGAAACTGTTCGCTCAG; R-GAGGTTGGTACATCAGAAACACC)

STAT1 (F-ATGGCAGTCTGGCGGCTGAATT; R-CCAAACCAGGCTGGCACAATTG)

STAT2 (F-CAGGTCACAGAGTTGCTACAGC; R-CGGTGAACTTGCTGCCAGTCTT)

STAT3 (F-CTTTGAGACCGAGGTGTATCACC; R-GGTCAGCATGTTGTACCACAGG)

OSMR (F-CAGGTGTTCCTACCAAATCTGCG; R-AATCCACCCTCTGTGCCTGCAA)

IL6ST (F-CACCCTGTATCACAGACTGGCA; R-TTCAGGGCTTCCTGGTCCATCA)

SOS1 (F-GGAGATCAACCCTTGAGTGCAG; R-TGCTCTACCCAGTGCCGACATA)

SOS2 (F-GGCATATCAGCAAACCAGGACAG; R-CACTCCCTACAAGTTCAGACGG)

PIK3CA (F-GAAGCACCTGAATAGGCAAGTCG; R-GAGCATCCATGAAATCTGGTCGC)

STAM2 (F-AGGTTGCACGGAAAGTGAGAGC; R-CCTCTGTGATTTTCTCCTTTCCAC)

CREBBP (F-AGTAACGGCACAGCCTCTCAGT; R-CCTGTCGATACAGTGCTTCTAGG)

CSF3R (F-CCACTACACCATCTTCTGGACC; R-GGTGGATGTGATACAGACTGGC)

SOX2-Qiagen RT2 #PPH02471A

NANOG (F-TGGGATTTACAGGCGTGAGCCAC; R-AAGCAAAGCCTCCCAATCCCAAAC)

OCT4 (F-GGGCTCTCCCATGCATTCAAAC; R-CACCTTCCCTCCAACCAGTTGC)

KLF4 (F-CGAACCCACACAGGTGAGAA; R-TACGGTAGTGCCTGGTCAGTTC)

NOTCH1 (F-CAATGTGGATGCCGCAGTTGTG; R-CAGCACCTTGGCGGTCTCGTA)

ASCL1 (F-CCCAAGCAAGTCAAGCGACA; R-AAGCCGCTGAAGTTGAGCC)

NSE-Qiagen RT2 #PPH02058A

SYP-Qiagen RT2 #PPH00717A

CHGA-Qiagen RT2 #PPH01181A

KRT5-Qiagen RT2 #PPH02625F

TP63-Qiagen RT2 #PPH01032F

KRT8-Qiagen RT2 #PPH02214F

KRT18-Qiagen RT2 #PPH00452F

CDH2-Qiagen QuantiTect #QT00063196

TGFb-Sigma-Aldrich #H_TGFB1_1

WNT5A-Qiagen QuantiTect #QT00025109

EPAS1-Qiagen RT2 #PPH02551C

SNAI1 (F-TGCCCTCAAGATGCACATCCGA; R-GGGACAGGAGAAGGGCTTCTC)

SMAD2 (F-GGGTTTTGAAGCCGTCTATCAGC; R-CCAACCACTGTAGAGGTCCATTC)

SMAD3 (F-TGAGGCTGTCTACCAGTTGACC; R-GTGAGGACCTTGTCAAGCCACT)

FN1 (F-ACAACACCGAGGTGACTGAGAC; R-GGACACAACGATGCTTCCTGAG)

TP53-Qiagen RT2 #PPH00213F

RB1-Qiagen RT2 #PPH00228F

Mouse qPCR primers for gene expression detection are listed:

Jak1 (F-CTGTCTACTCCATGAGCCAGCT; R-CCTCATCCTTGTAGTCCAGCAG)

Stat3 (F-AGGAGTCTAACAACGGCAGCCT; R-GTGGTACACCTCAGTCTCGAAG)

Tp53 (F-TGAAGGCCCAAGTGAAGCCCTC; R-TGTGGCGCTGACCCACAACTGC)

Rb1 (F-CCTTGAACCTGCTTGTCCTCTC; CTGAGGCTGCTTGTGTCTCTGT)

Stat1 (F-GCCTCTCATTGTCACCGAAGAAC; R-TGGCTGACGTTGGAGATCACCA)

Stat2 (F-GAACCAACTCTCCATTGCCTGG; R-CGTAAGAGGAGAACTGCCAGCT)

Sox2 (F-AACGGCAGCTACAGCATGATGC; R-CGAGCTGGTCATGGAGTTGTAC)

Krt5 Qiagen RT2 #PPM59967F-200

Trp63 Qiagen RT2 #PPM03458A-200

Krt8 Qiagen RT2 #PPM04776F-200

Krt18 Qiagen RT2 #PPM05184A-200

OCT4 Qiagen RT2 #PPM68766A-200

CDH2 Qiagen QuantiTect #QT00148106

SNAI1 Qiagen QuantiTect #QT00240940

SNAI2 Qiagen QuantiTect #QT00098273

EPAS1 (F-GGACAGCAAGACTTTCCTGAGC; R-GGTAGAACTCATAGGCAGAGCG)

TGFB (F-TGATACGCCTGAGTGGCTGTCT; R-CACAAGAGCAGTGAGCGCTGAA)

KLF4 (F-GAACGCCTCATCAATGCCTGCA; R-GAATCAGGGCTGCCTTGAAGAG)

#### ChIP-qPCR and SOX2 ChIP-seq

ChIP experiments were performed as previously described (*5, 10*). Briefly, cultured cells were crosslinked with 1% formaldehyde and was quenched with 0.125M glycine. Cells were then rinsed with cold 1X PBS twice and lysed in 1% SDS containing buffer supplemented with 1X protease and phosphatase inhibitors. Chromatin was sonicated to an average length of 500bp and then centrifuged at 14,000 rpm to remove the debris. One percent of the supernatant was saved as input, and the rest was added with ChIP-grade antibody overnight, then added 20ul of agarose/protein A or G beads and incubated for 4 hours. Beads were washed with standard wash buffers (Low-Salt, High-Salt, and LiCl) and finally with TE. The immunoprecipitated chromatin were eluted in elution buffer and de-crosslinked by NaCl at 65°C overnight. Proteins were then digested by proteinase K and DNA was purified with MinElute PCR Purification Kit (Qiagen, Cat #28006) and eluted with 10ul water. Antibodies used are Anti-Histone H3 (acetyl K27) antibody-ChIP Grade (Abcam, cat# ab4729), Anti-Histone H3 (tri methyl K4) antibody-ChIP Grade(Abcam,cat# ab8580), Tri-Methyl-Histone H3 (Lys27) (C36B11) Rabbit mAb (Cell Signaling Technology, cat #9733S). ChIP-qPCR primers:

Jak1-1-F: TGCTTCCCTCCCAAATACACCTCA;

Jak1-1-R: TTCCTGCTTTGCACTTCAGCTCAG (H3K27me3);

Jak1-2-F: GTGAATGGTCCATCCCCACA;

Jak1-2-R: TTTCCCAAAGTGGGGCACAA (H3K27ac/ H3K4me3).

SOX2 ChIP-Seq in the CWR-R1 and WA01 cells were described in Larischa et al., in revision. ChIP experiments were conducted using the ChIP Assay Kit per the manufacturer’s protocol (EMD Millipore; Burlington, MA). A polyclonal goat anti-SOX2 mAb (P48431, R&D Systems; Minneapolis, MN) or goat IgG control were used for immunoprecipitation. Eluted ChIP DNA was purified using the PCR Purification Kit (Qiagen). ChIP-Seq libraries were generated using the KAPA LTP Library Preparation Kit (#KK8230; Kapa Biosystems; Wilmington, MA). Libraries were sequenced on a HiSeq 2000 sequencing system (Illumina) in a 50-bp, single-end run.

#### Bulk RNA-seq preparation and analysis

LNCaP/AR cells transduced with different CRISPR constructs were treated with enzalutamide or vehicle for 6 days before the total RNA was extracted using Trizol (Ambion, Cat 15596018) as previously described (*7, 10*). RNA-Seq libraries were prepared using the Illumina TruSeq stranded mRNA kit, with 10 cycles of PCR amplification, starting from 500 ng of total RNA, at the integrated genomics operation (IGO) Core at MSKCC. Barcoded RNA-Seq were run as paired-end read 50 nucleotides in length on the Illumina HiSeq 2500 and Poly-A selection was performed. Adapter trimming and quality trimming was performed with trimgalore (v0.5.0), and ribosomal RNA was removed using SortMeRNA (v4.1.0). Trimmed and filtered reads were aligned to reference (GRCh37) with STAR (vSTAR2.6.1d). FeatureCounts (v1.6.4) was used for gene counts, biotype counts, and rRNA estimation. FPKMs for genes and transcripts were generated by StringTie (v1.3.5), and RSeQC (v3.0.0) was used for generating RNA quality control metrics. Differential gene expression analysis was performed using the R package DEseq2 (v1.6.3). Cutoff values of absolute fold change greater than 2 and FDR<0.1 were used to select for differentially expressed genes between sample group comparisons. GSEA statistical analysis was carried out with the R package fgsea (https://www.biorxiv.org/content/10.1101/060012v3).

#### Single cell RNA-seq preparation and analysis

LNCaP/AR cells transduced with different CRISPR constructs were treated with enzalutamide or vehicle for 5 days before the cells were collected. Single-cell RNA-seq were performed by the 10x Genomic single cell 5’library platform. Based on FACS analysis, single cells were sorted into 1.5 ml tubes (Eppendorf) and counted manually under the microscope. The concentration of single cell suspensions was adjusted to 900-1100 cells/μl. Cells were loaded between 10,000 and 17,000 cells/chip position using the Chromium Single cell 5’ Library, Gel Bead & Multiplex Kit and Chip Kit (10x Genomics, V1 barcoding chemistry). Single-cell gene expression libraries were generated according to the manufacturer’s instructions and single-cell expression sequencing was run on a NovaSeq 6000 (Novogene Co., Ltd). All the subsequent steps were performed following the standard manufacturer’s protocols. 10x scRNA-seq data was preprocessed using the Cell Ranger software (5.0.0). We used the “mkfastq”, “count” and ‘aggr’ commands to process the 10x scRNA-seq output into one cell by gene expression count matrix, using default parameters. scRNA-seq data analysis was performed with the Scanpy (1.6.0) package in Python(*68*). Genes expressed in fewer than 3 cells were removed from further analysis. Cells expressing less than 100 and more than 7000 genes were also removed from further analysis. In addition, cells with a high (>= 0.15) mitochondrial genome transcript ratio were removed. For downstream analysis, we used count per million normalization (CPM) to control for library size difference in cells and transformed those into log(CPM+1) values. After normalization, we used the ‘pp.highly_variable_genes’ command in Scanpy to find highly variable genes across all cells using default parameters except for “min_mean = 0.01”. The data were then z-score normalized for each gene across all cells. We then used the ‘tl.pca (n_comps=50, use_highly_variable=True)’, the ‘pp.neighbors (n_pcs=25, n_neighbors=15)’ and the ‘tl.leiden (resolution = 0.75)’ command in Scanpy to partition the single cells into 6 distance clusters. Briefly, these processes first identify 50 principal components in the data based on the previously found highly variable genes to reduce the dimensions in the original data, and then build a nearest neighbor graph based on the top 25 principal components, and finally a partition of the graph that maximizes modularity was found with the Leiden algorithm(*69*). To evaluate the activity of lineage specific transcriptional programs in those cells, we utilized a custom library of genes based on the well-established gene signatures for AR target genes (AR score) and NE, luminal, basal, stem-like and EMT lineages. The AR score gene signature was adapted from Hieronymus et al(*48*), luminal, basal and NE gene signatures were defined by combining the signature genes from(*10, 11, 46, 47*). EMT and stem-like gene signature were adapted from the signature genes of Dong et al(*47*)plus canonical lineage marker genes (table S2). The activation score was calculated based on the overall expression of genes in each gene list using the ‘tl.score_genes’ function of the Scanpy package.

## Statistics Methods

All of the statistical details of experiments can be found in figure legends. For comparisons between two groups of independent datasets when normality and homoscedasticity are satisfied, multiple t tests were performed, p value and standard error of the mean (s.e.m.) were reported. For comparing gene expressions between two patients’ groups, Mann Whitney U Test (Wilcoxon Rank Sum Test) were performed. For comparisons among more than two groups (>2), one-way or two-way ANOVA were performed, p values and s.e.m. were reported; and p values were adjusted by multiple testing corrections when applicable. For dose response curve, p values were calculated by non-linear regression with extra sum-of-squares F test. Fisher’s Exact test was used to compare the frequency of genomic alterations between different patients’ group and percentage of cell populations. Chi-square test with Yates correction were used to compared the exact cell numbers of different clusters of single cell subclones. For all figures, **** represents p<0.0001. *** represents p<0.001. ** represents p<0.01. * represents p<0.05.

---

**Fig.S1:**
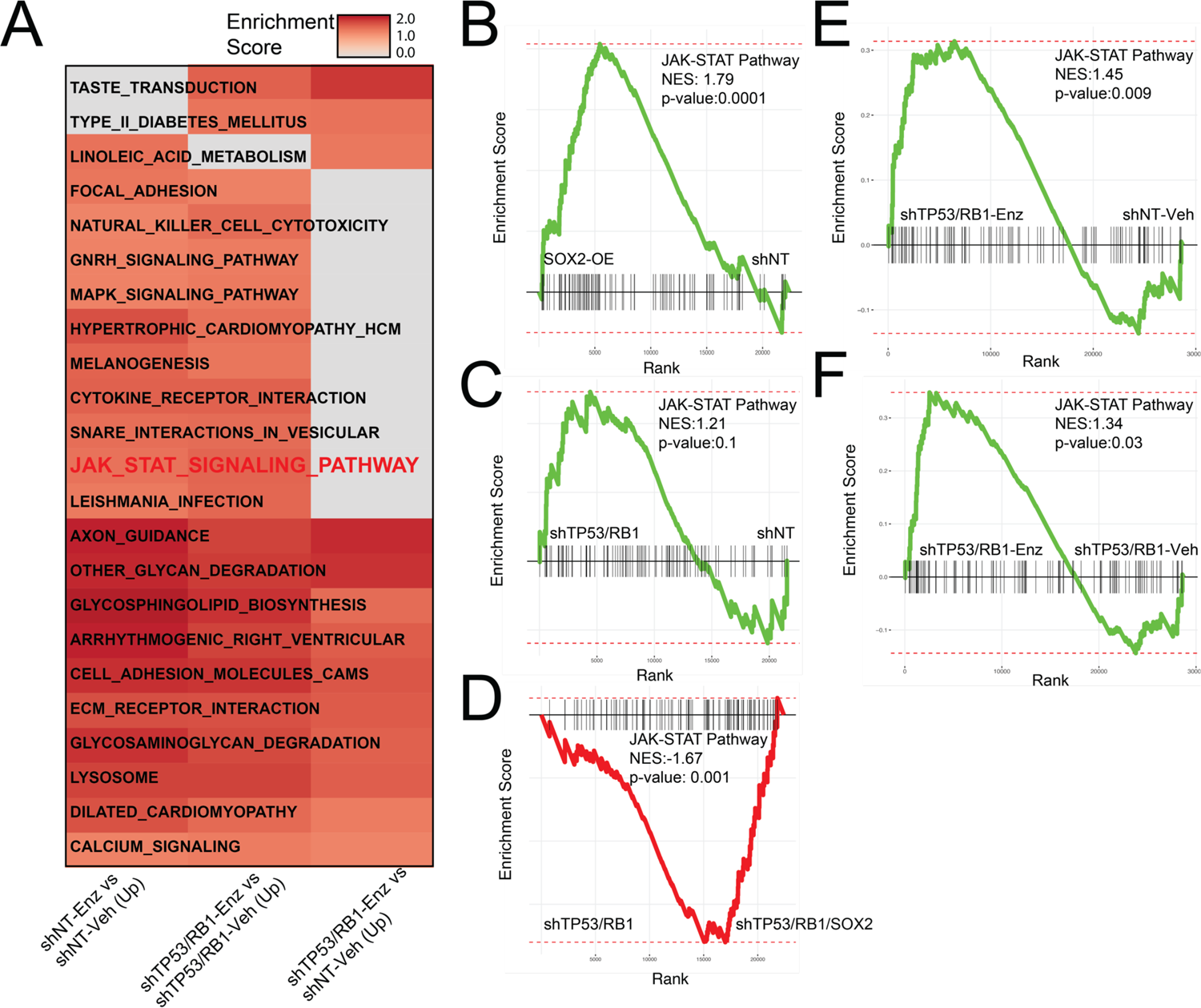
JAK-STAT signaling pathway is enriched in enzalutamide resistant mCRPC with TP53/RB1-deficiency. (**A**) Heatmap represents the significant changed signaling pathways in LNCaP/AR cell lines transduced with annotated shRNAs and treated with enzalutamide or vehicle, based on GSEA analysis. Three comparations are presented and reads from 3 biological replicates in each group were used for analysis. (**B-F**) GSEA analysis of JAK-STAT signaling pathway (KEGG_JAK_STAT_Signaling_Pathway) expression in: (**B**) SOX2-OE group compared to shNT group; (**C**) shTP53/RB1 group compared to shNT group; (**D**) shTP53/RB1 group compared to shTP53/RB1/SOX2 group; (**E**) shTP53/RB1+Enz group compared to shNT-Veh group; (**F**) shTP53/RB1+Enz group compared to shTP53/RB1+Veh group. Reads from 3 biological replicates were used for analysis.

**Fig. S2.**
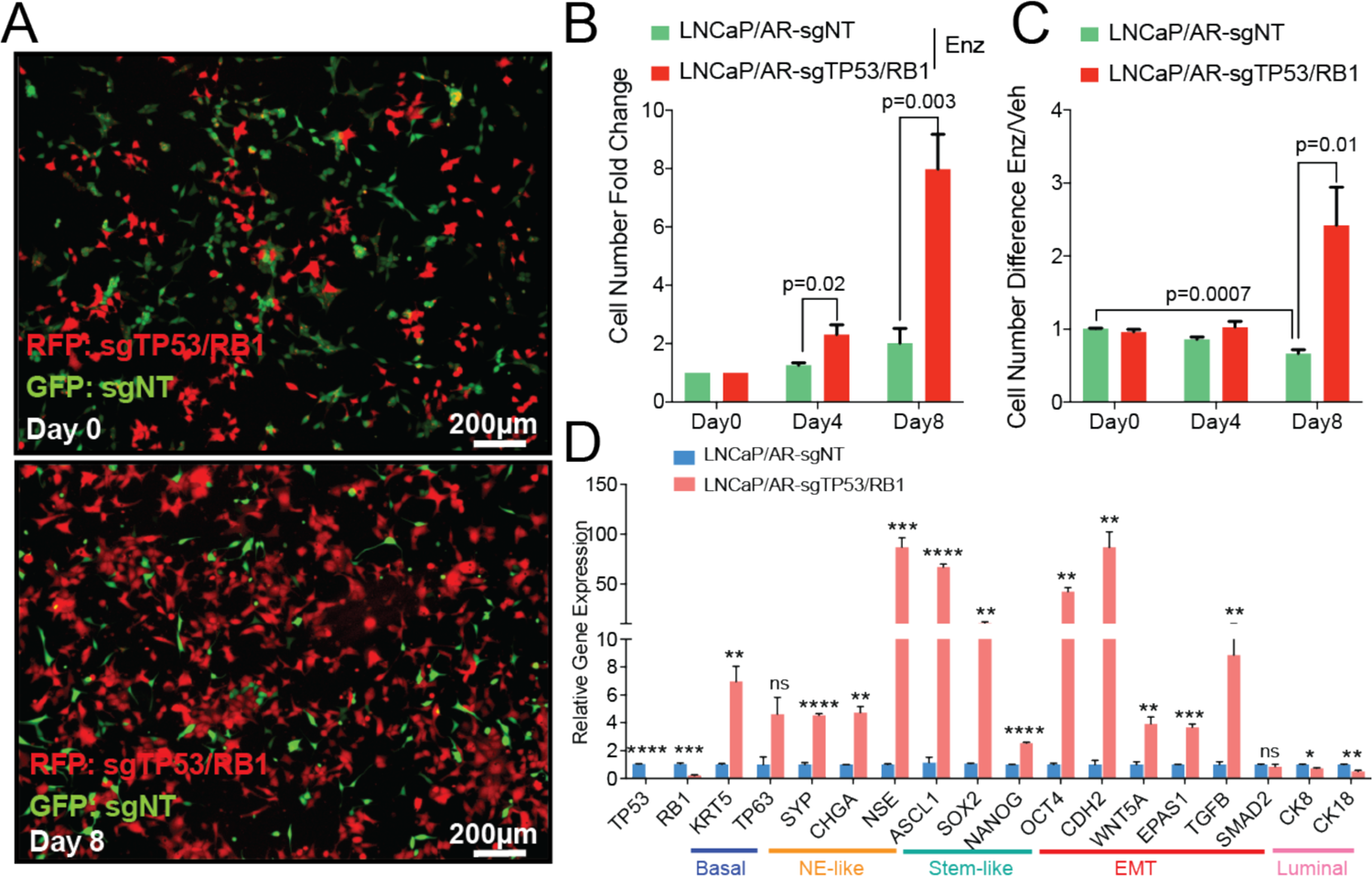
LNCaP/AR-sgTP53/RB1 is a highly resistant and lineage plastic cell line model. (**A**) Fluorescence microscope imaging shows the cell mixtures of sgTP53/RB1-RFP cells (red) and sgNT-GFP cells (green) on Day 0 and Day 8 of the competition assay cultured in CSS medium and 10µM enzalutamide. (**B**) Relative cell number of LNCaP/AR cells transduced with annotated guide RNAs measured in the competition assay. (**C**) Cell number fold change of LNCaP/AR cells transduced with annotated guide RNAs. For **B-C**, cells were treated with 10 μM enzalutamide (Enz) or DMSO (Veh) for 8 days in CSS medium and cell number was measured using FACS. (**D**) Relative expression of canonical lineage marker genes in LNCaP/AR mCRPC cells transduced with annotated guide RNAs. For all panels, mean ± s.e.m. is represented, and p values were calculated using multiple t tests. **** p<0.0001. *** p<0.001. ** p<0.01. * p<0.05.

**Fig. S3.**
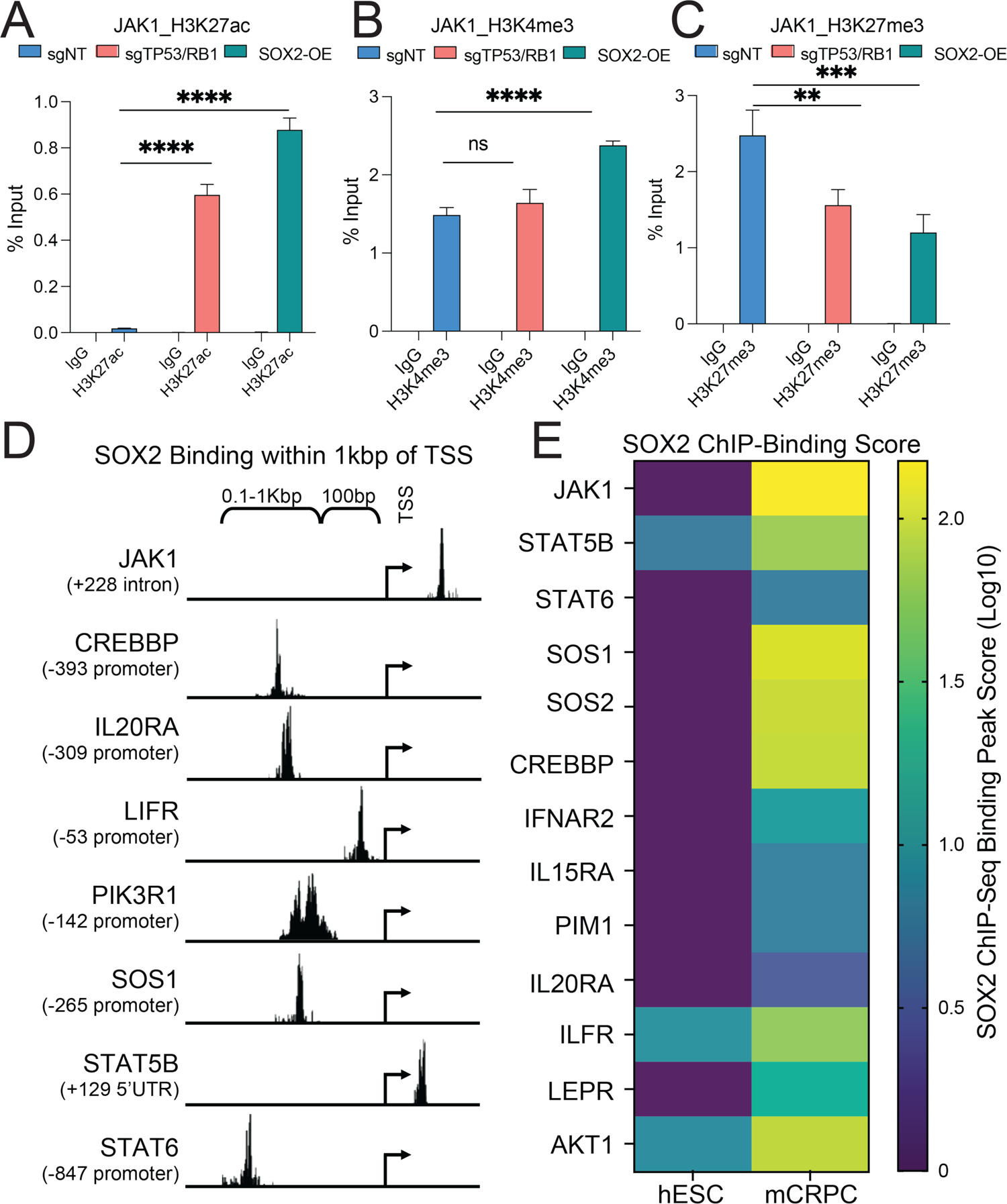
TP53/RB1-deficiency and Sox2 overexpression promotes the transcriptional activation of JAK1. (**A**) H3K27ac ChIP-qPCR of the *JAK1* genomic locus in LNCaP/AR cells transduced with annotated constructs. (**B**) H3K4me3 ChIP-qPCR of the *JAK1* genomic locus in LNCaP/AR cells transduced with annotated constructs. (**C**) H3K27me3 ChIP-qPCR of the *JAK1* genomic locus in LNCaP/AR cells transduced with annotated constructs. For all panels unless otherwise noted, mean ± s.e.m. is represented and p values were calculated using one-way ANOVA. (**D**) Representative SOX2 binding sites in the genomic loci of JAK-STAT signaling genes in the mCRPC CWR-R1 cell line based on ChIP-seq analysis. (**E**) SOX2 binding peak score in the genomic loci of JAK-STAT signaling genes in the mCRPC CWR-R1 cell (prostate cancer specific binding) compared to human ESC cell line WA01.

**Fig. S4.**
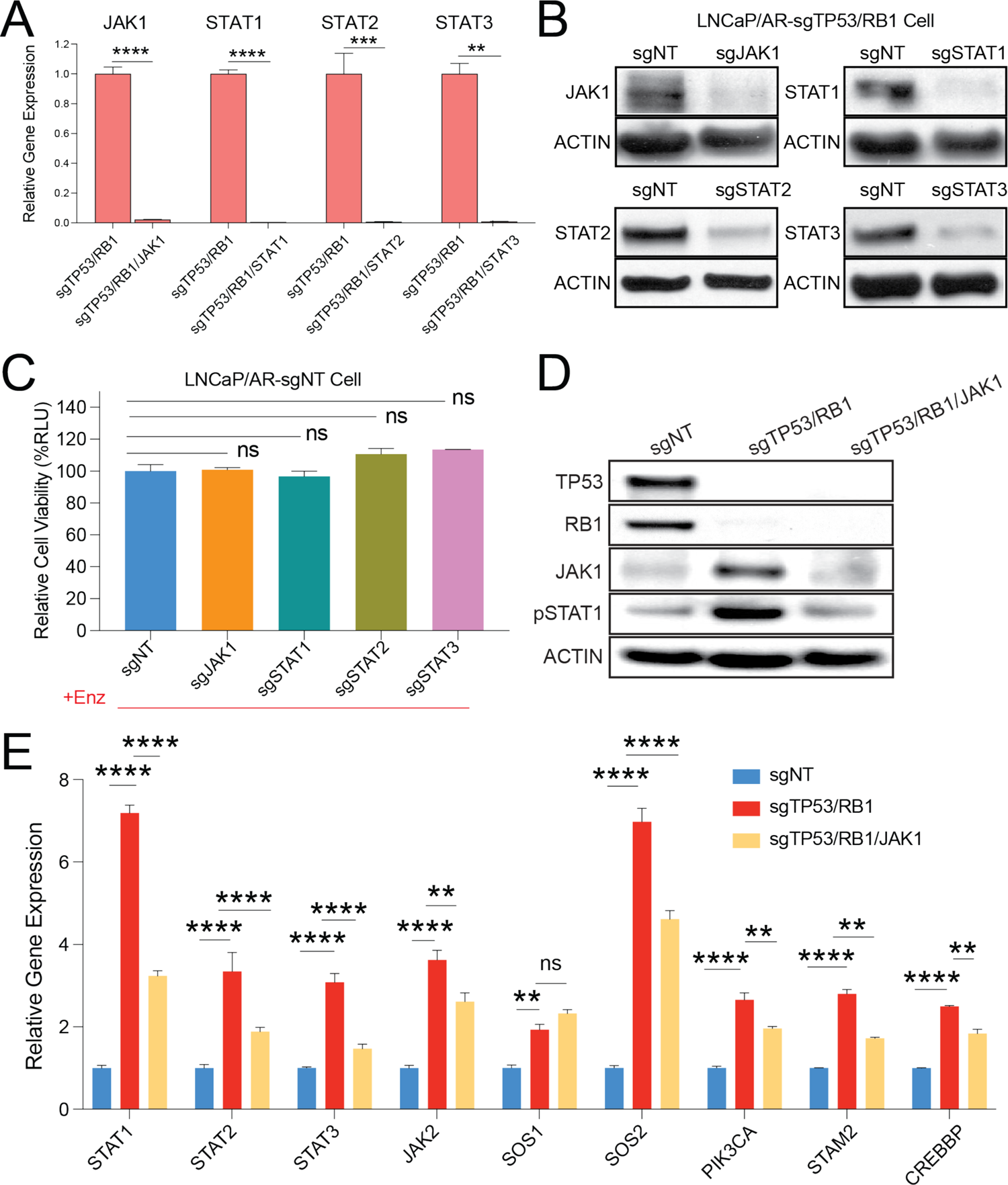
JAK-STAT signaling is significantly impaired in the sgTP53/RB1/JAK1 cells. (**A**) Relative expression of JAK-STAT genes in LNCaP/AR-sgTP53/RB1 cells transduced with annotated guide RNAs. (**B**) Western blot of JAK1, STAT1-3 proteins in LNCaP/AR cells transduced with annotated guide RNAs. (**C**) Relative cell number of LNCaP/AR-sgNT cells transduced with annotated CRISPR guide RNAs. Cells were treated with 10 µM enzalutamide (Enz) for 8 days and cell number was measured using CellTiter-Glo assay, all normalized to sgNT group. (**D**) Western blot of JAK1 and pSTAT1 proteins in LNCaP/AR cells transduced with annotated guide RNAs. (**E**) Relative expression of canonical JAK-STAT genes in LNCaP/AR mCRPC cells transduced with annotated guide RNAs. p values were calculated using two-way ANOVA. mean ± s.e.m. is represented and **** p<0.0001. *** p<0.001. ** p<0.01. * p<0.05. **** p<0.0001. *** p<0.001. ** p<0.01. * p<0.05.

**Fig. S5.**
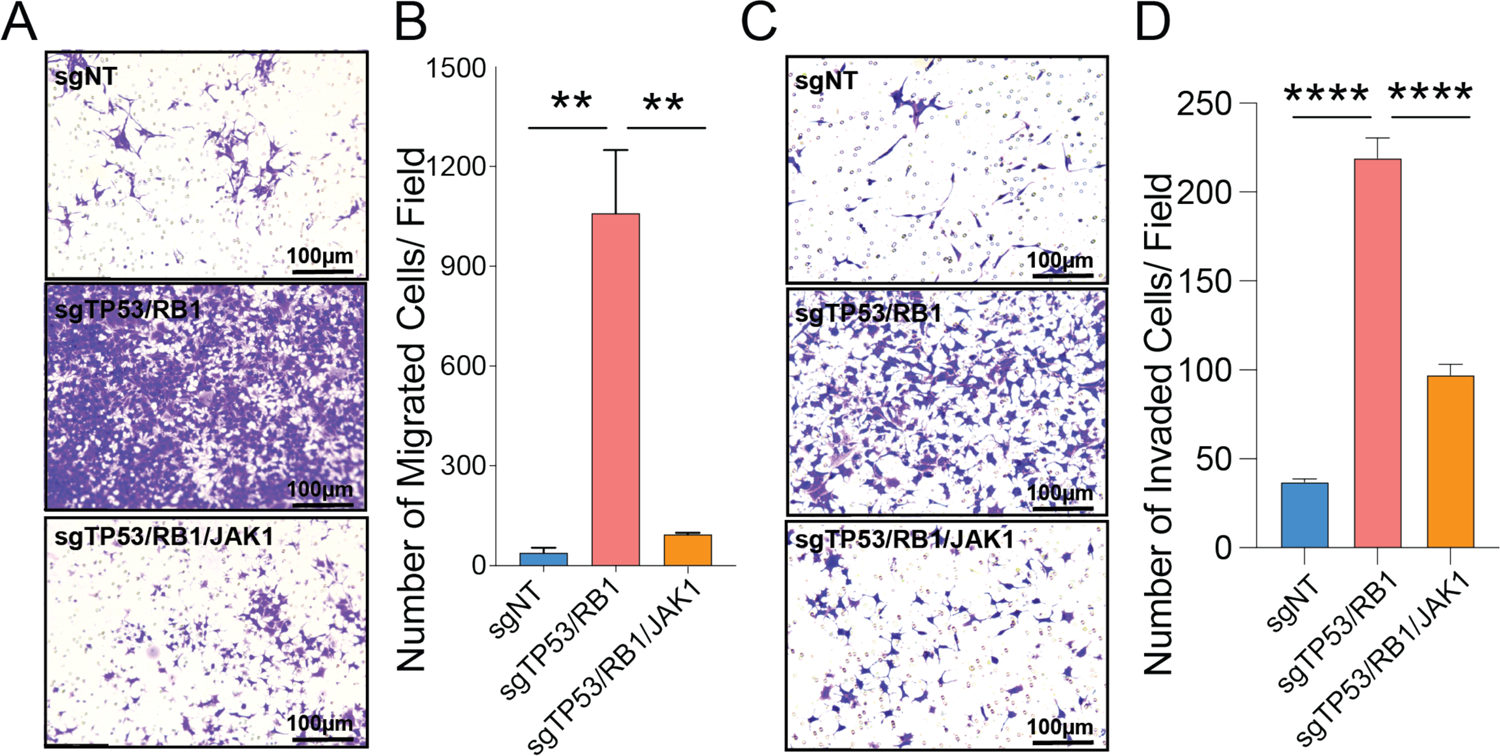
JAK1-KO significantly reversed the increased migration and invasion ability of mCRPC cells with TP53/RB1-deficiency. (**A**) Representative pictures of LNCaP/AR cell (sgNT, sgTP53/RB1, sgTP53/RB1/JAK1) transwell migration assay. 9 representative pictures were taken for each cell line and scale bar is annotated. (**B**) Quantification of the migrated cell numbers of 9 representative images for each of the cell lines. (**C**) Representative pictures of LNCaP/AR cell (sgNT, sgTP53/RB1, sgTP53/RB1/JAK1) invasion assay. 9 representative pictures were taken for each cell line and scale bar is annotated. (**D**) Quantification of the invaded cell numbers of 9 representative images for each of the cell lines. For all panels, p value was calculated by one way ANOVA. mean ± s.e.m. is represented and **** p<0.0001. *** p<0.001. ** p<0.01. * p<0.05. **** p<0.0001. *** p<0.001. ** p<0.01. * p<0.05.

**Fig. S6.**
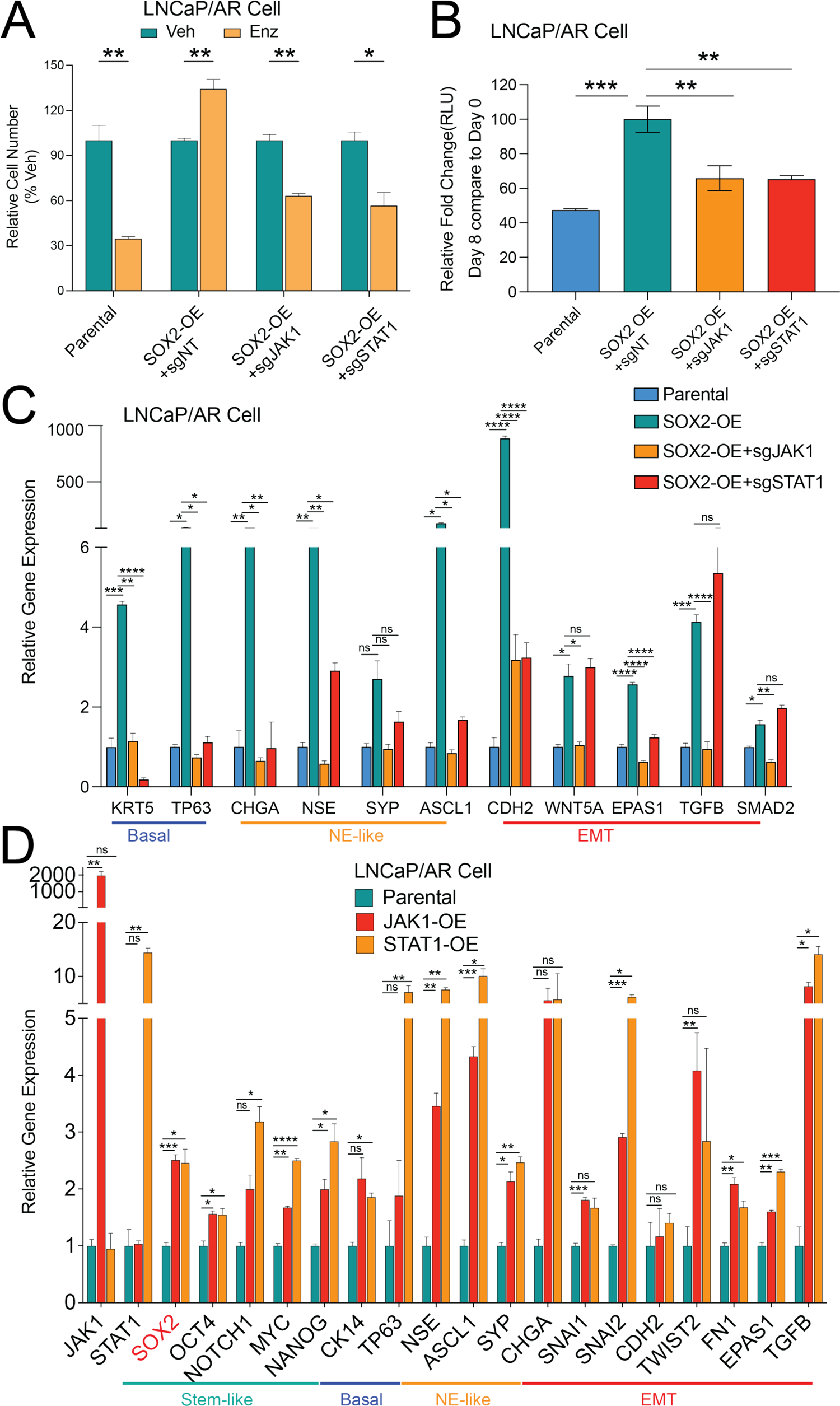
JAK-STAT signaling is required for the SOX2-mediated lineage plasticity and resistance. (**A**) Relative cell number of LNCaP/AR cells transduced with annotated constructs and treated with various treatments, normalized to “Veh” group. Enz denotes 10μM enzalutamide, Veh denotes DMSO treatment with same volume as enzalutamide, for 6 days and cell number were measured by cell proliferation assay. mean ± s.e.m. is represented, and p values were calculated using 2-tailed multiple t-test. (**B**) Relative cell number fold change of LNCaP/AR cells transduced with annotated constructs and treated with various treatments, normalized to “Veh” group. Enz denotes 10μM enzalutamide, Veh denotes DMSO treatment with same volume as enzalutamide, for 6 days and cell number were measured by CellTiterGlo assay. mean ± s.e.m. is represented, and p values were calculated using one-way ANOVA. (**C**) Relative expression of canonical lineage marker genes in LNCaP/AR-SOX2-OE cells transduced with annotated constructs. Mean ± s.e.m. is represented, and p values were calculated using 2-way ANOVA. (**D**) Relative expression of canonical lineage marker genes in LNCaP/AR cells transduced with JAK1 or STAT1 cDNA constructs. SOX2 expression is highlighted in red. mean ± s.e.m. is represented, and p values were calculated using two-way ANOVA. For all panels, **** p<0.0001. *** p<0.001. ** p<0.01. * p<0.05. **** p<0.0001. *** p<0.001. ** p<0.01. * p<0.05.

**Fig. S7.**
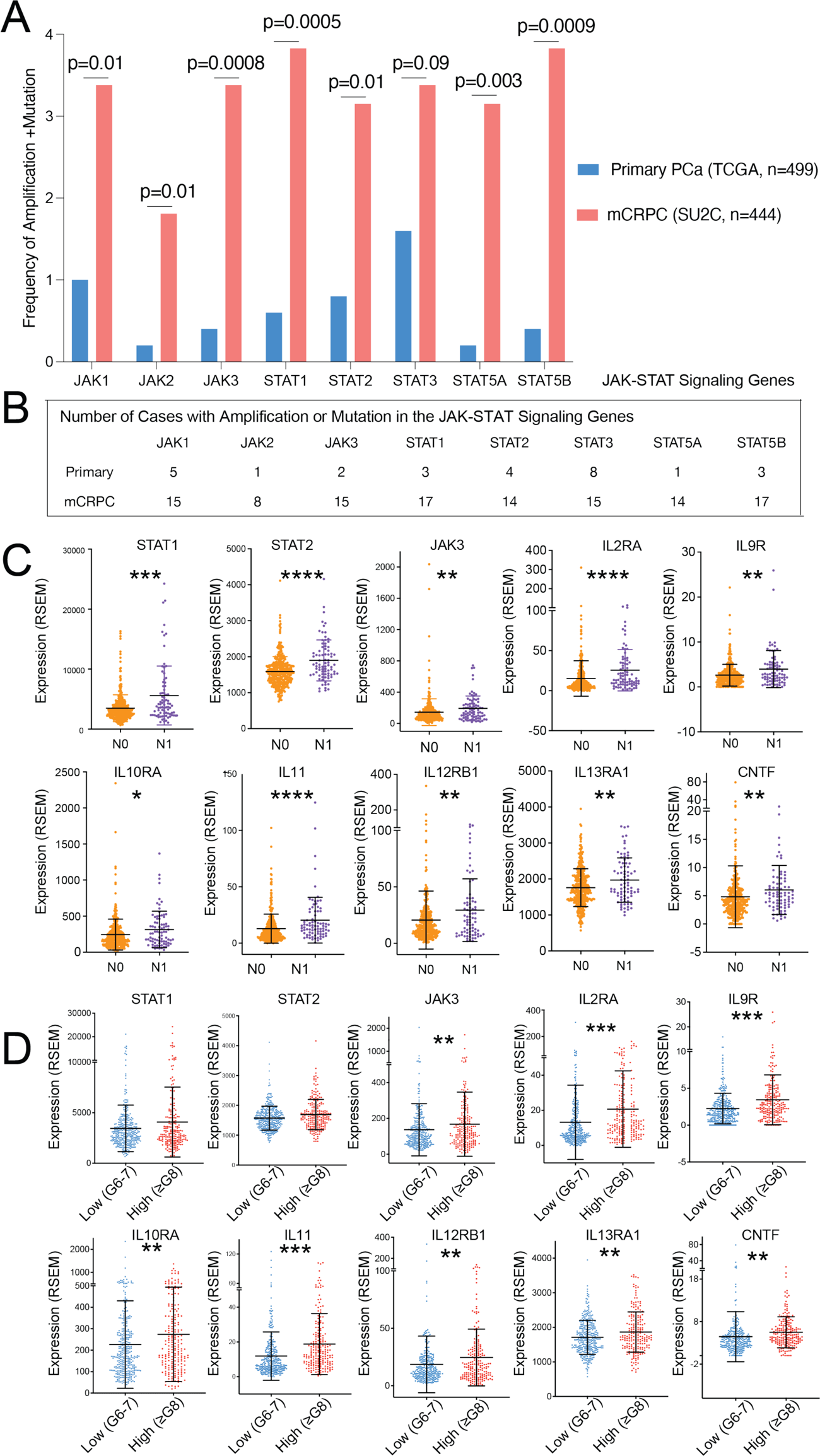
JAK1 and STAT1 genomic alterations is correlated with poor outcome of patients with mCRPC. (**A**) Frequency of amplification or mutations in the genomic loci of key JAK-STAT signaling genes in the mCRPC tumors of the SU2C cohort, compared to the frequency in the primary tumors of the TCGA cohort. p values were calculated using two-tails Fisher’s exact test. (**B**) Number of cases with amplification or mutations in the genomic loci of key JAK-STAT signaling genes, in the SU2C cohort, compared to the TCGA cohort. (**C**) Expression (RSEM) of JAK-STAT signaling genes in patients with regional lymph nodes metastasis (N1, n=80) compared to the ones without regional lymph nodes metastasis (N0, n=345). (**D**) Expression (RSEM) of JAK-STAT signaling genes in the high-grade tumors (Gleason score ≥8, n=206) compared to the low-grade tumors (Gleason score≤7, n=292). For panel **C-D**, mean ± s.d. is represented and p values were calculated using Mann-Whitney test. **** p<0.0001. *** p<0.001. ** p<0.01. * p<0.05. **** p<0.0001. *** p<0.001. ** p<0.01. * p<0.05.

**Fig. S8.**
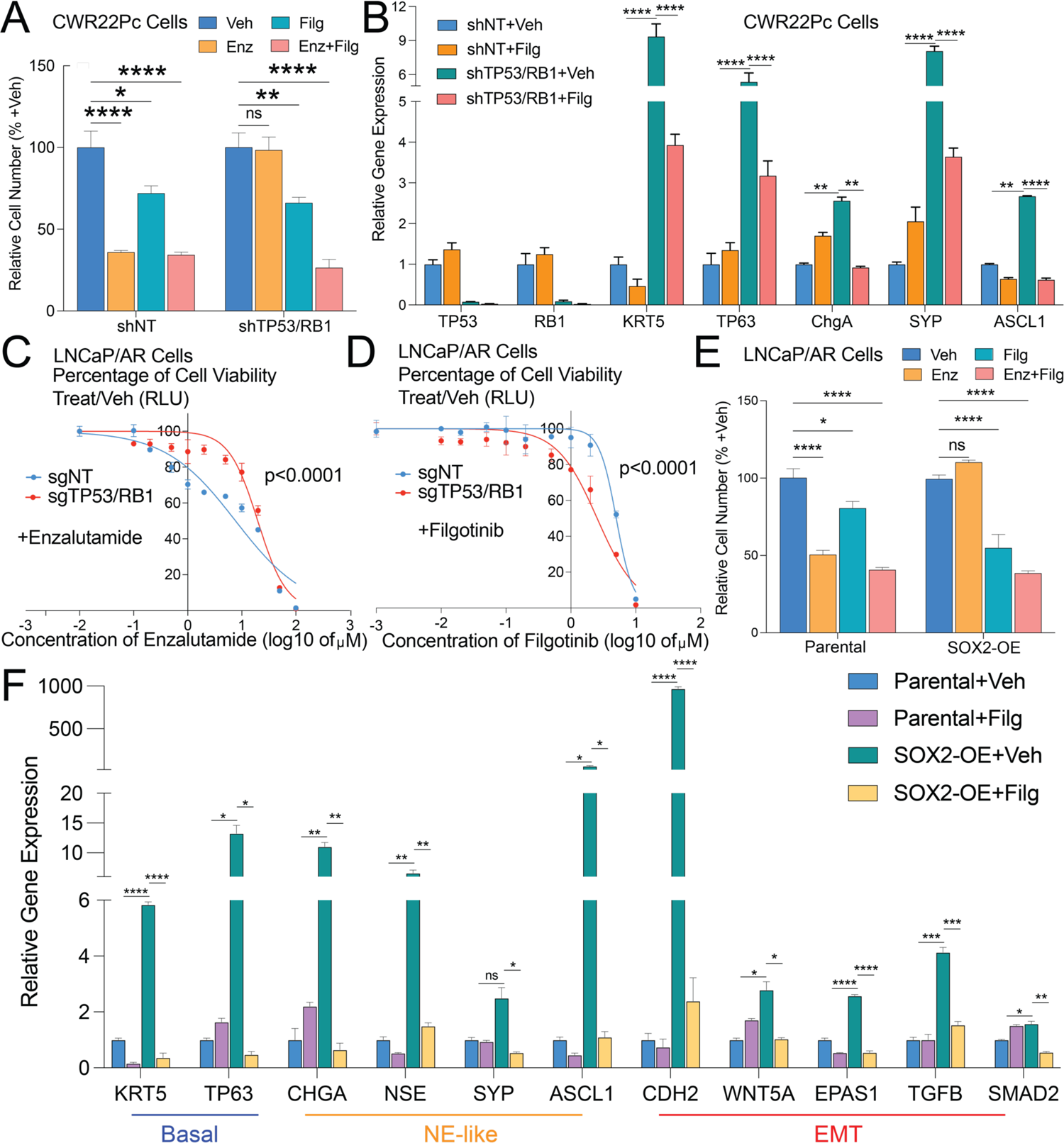
JAK inhibitor impairs lineage plasticity and restore enzalutamide sensitivity. (**A**) Relative cell number of CWR22Pc cells transduced with annotated CRISPR guide RNAs and treated with various treatments, normalized to “Veh” group. Enz denotes 1μM enzalutamide, Filg denotes 5μM filgotinib, Enz+Filg denotes the combination of enzalutamide and filgotinib, Veh denotes DMSO treatment with same volume as enzalutamide. Cells were treated for 4 days and cell number were counted. (**B**) Relative expression of canonical lineage marker genes in CWR22Pc cells transduced with annotated shRNAs and treated with vehicle or filgotinib, normalized to “shNT+Veh” group. Filg denotes 5μM filgotinib, Veh denotes DMSO treatment with same volume as filgotinib. (**C**) Enzalutamide dose response curve of LNCaP/AR cells transduced with annotated CRISPR guide RNAs. p values were calculated by non-linear regression with extra sun-of-squares F test, 3 biological replicates were used for each data point. (**D**) Filgotinib dose response curve of LNCaP/AR cells transduced with annotated CRISPR guide RNAs. p values were calculated by non-linear regression with extra sun-of-squares F test, 3 biological replicates were used for each data point. (**E**) Relative cell number of LNCaP/AR cells transduced with annotated constructs and treated with various treatments for 8 days, normalized to “Veh” group. Enz denotes 10μM enzalutamide, Filg denotes 10μM filgotinib, Enz+Filg denotes the combination of enzalutamide and filgotinib, Veh denotes DMSO treatment with same volume as enzalutamide, for 8 days and cell number were counted. (**F**) Relative expression of canonical lineage marker genes in LNCaP/AR-SOX2-OE cells treated with annotated treatments. Enz denotes 10μM enzalutamide, Veh denotes DMSO treatment with same volume as enzalutamide. Cells were treated for 6 days. mean ± s.e.m. is represented, and p values were calculated using two-way ANOVA. For all panels, ***** p<0.0001. *** p<0.001. ** p<0.01. * p<0.05.

**Fig. S9.**
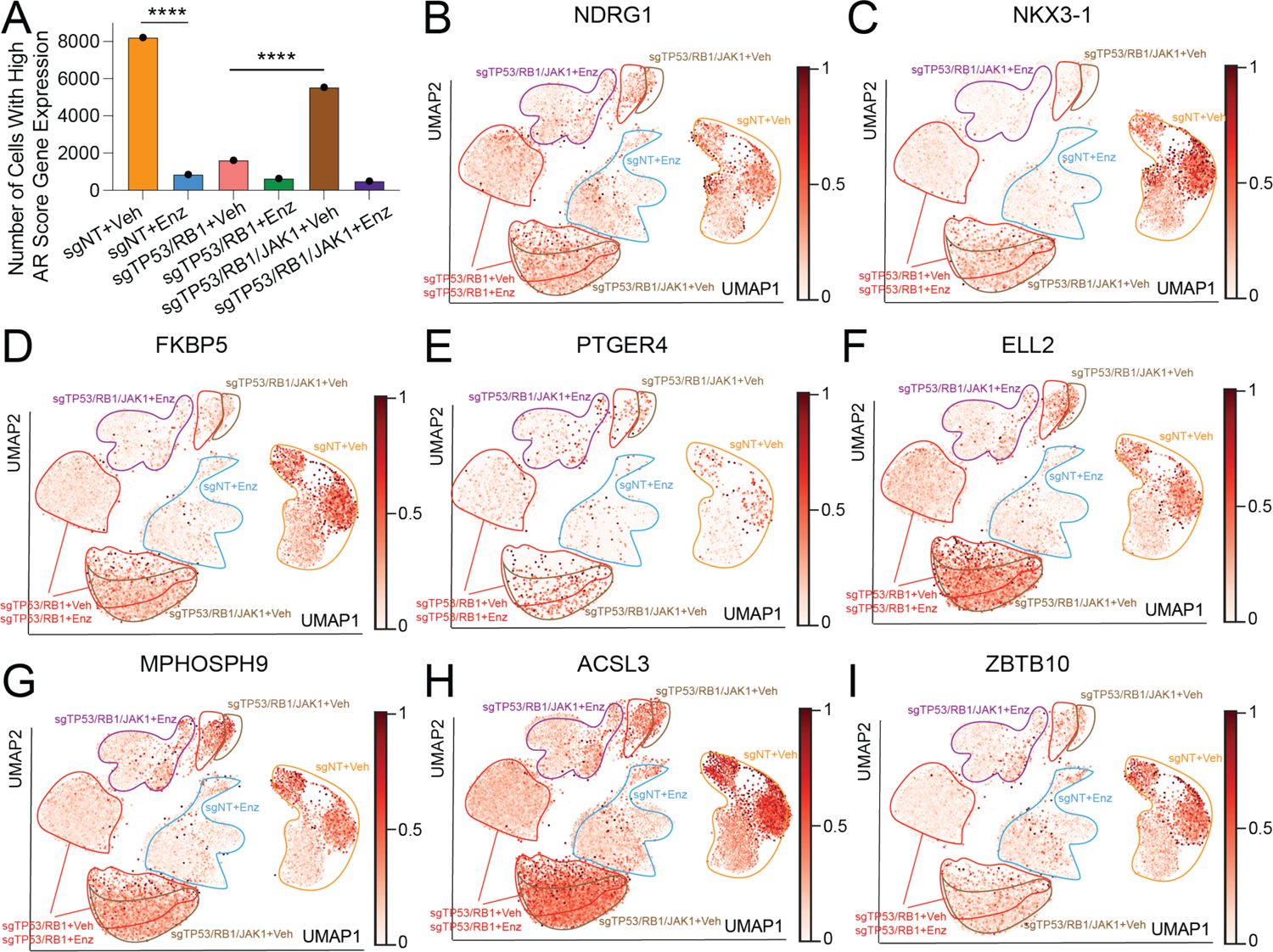
AR signaling partially restored in the subclones with TP53/RB1/JAK1-KD and vehicle treatment. (**A**) Bar plot presents the number of single cells express high level (expression level in the top 20% of all single cells of all samples) of AR targeted genes (partial AR Score genes as shown in table S2). p-values are calculated with Chi-square test with Yates correction. **** p<0.0001, *** p<0.001,** p<0.01, * p<0.05. ns: not significant. (**B-I**) UMAP plot of single cell transcriptomic profiles colored by expression of selected AR target genes (z-score, AR Score genes) for each cell (dot) of LNCaP/AR cells transduced with annotated CRISPR guide RNAs and treated with vehicle or enzalutamide for 5 days. Color density of each cell is scaled by the color bar. Fields of different sample groups are labeled with different color.

**Fig. S10.**
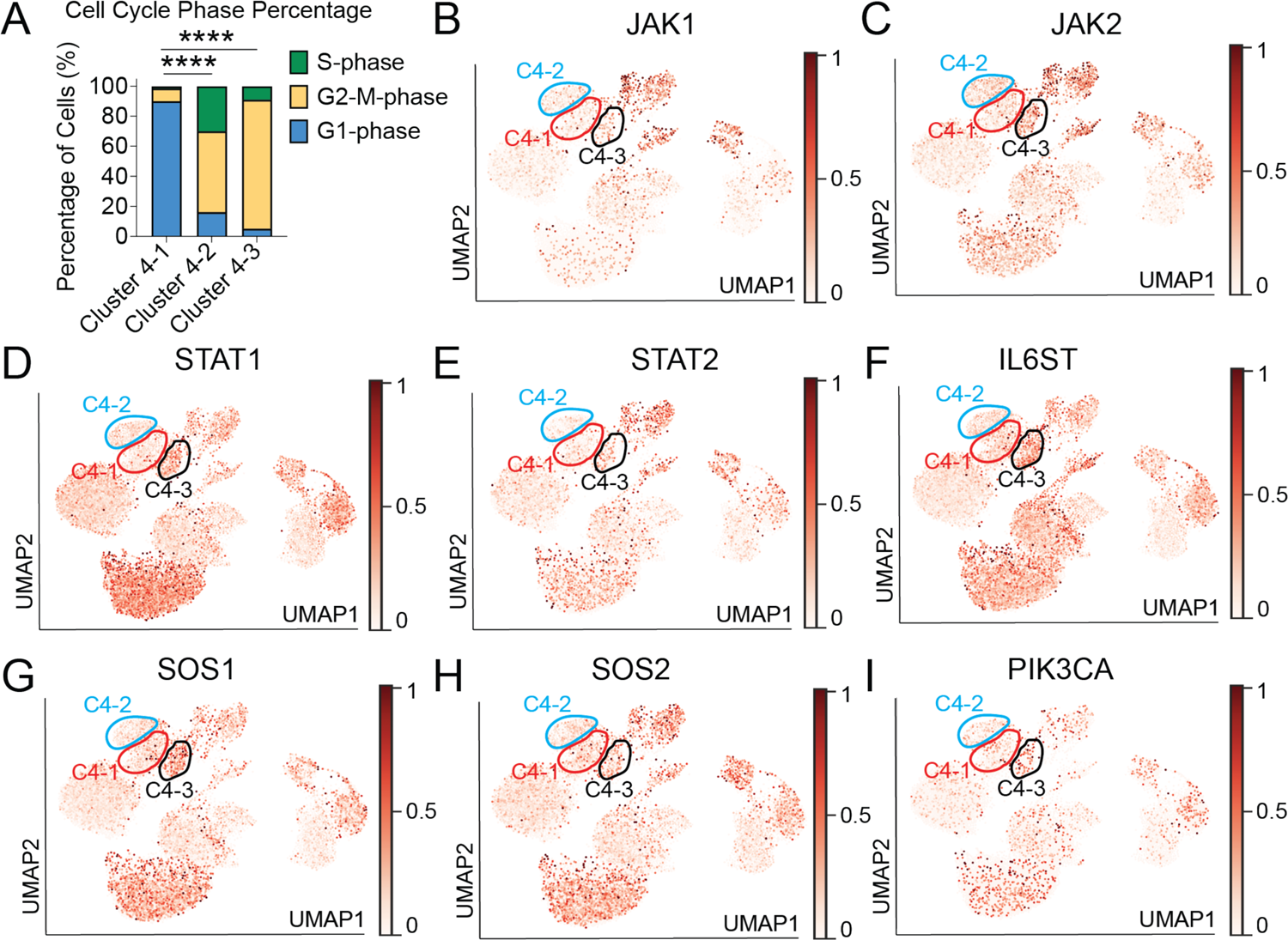
Subclusters within the Cluster 4 display remaining and various levels of JAK-STAT signaling: **(A)** Bar plot presents the percentage distribution of each single cells in different cell cycle phases in subcluster 4-1, 4-2 and 4-3. p-values are calculated with Fisher’s Exact Test. **** p<0.0001. (**B-I**) UMAP plot of single cell transcriptomic profiles colored by expression of canonical JAK-STAT target genes (z-score) for each cell (dot) of LNCaP/AR cells transduced with annotated CRISPR guide RNAs and treated with vehicle or enzalutamide for 5 days. For panel **B-I**, distribution area of subcluster 4-1, 4-2, 4-3 are labeled with red, blue and black. Color density of each cell is scaled by the color bar.

**Table S1: Gene Set Enrichment Analysis (GSEA) results show significantly changed signaling pathways.** GSEA results of six different comparations, including the enriched gene lists, are presented in attached excel file: table S1_GSEA results.xlsx.

**TableS2:**
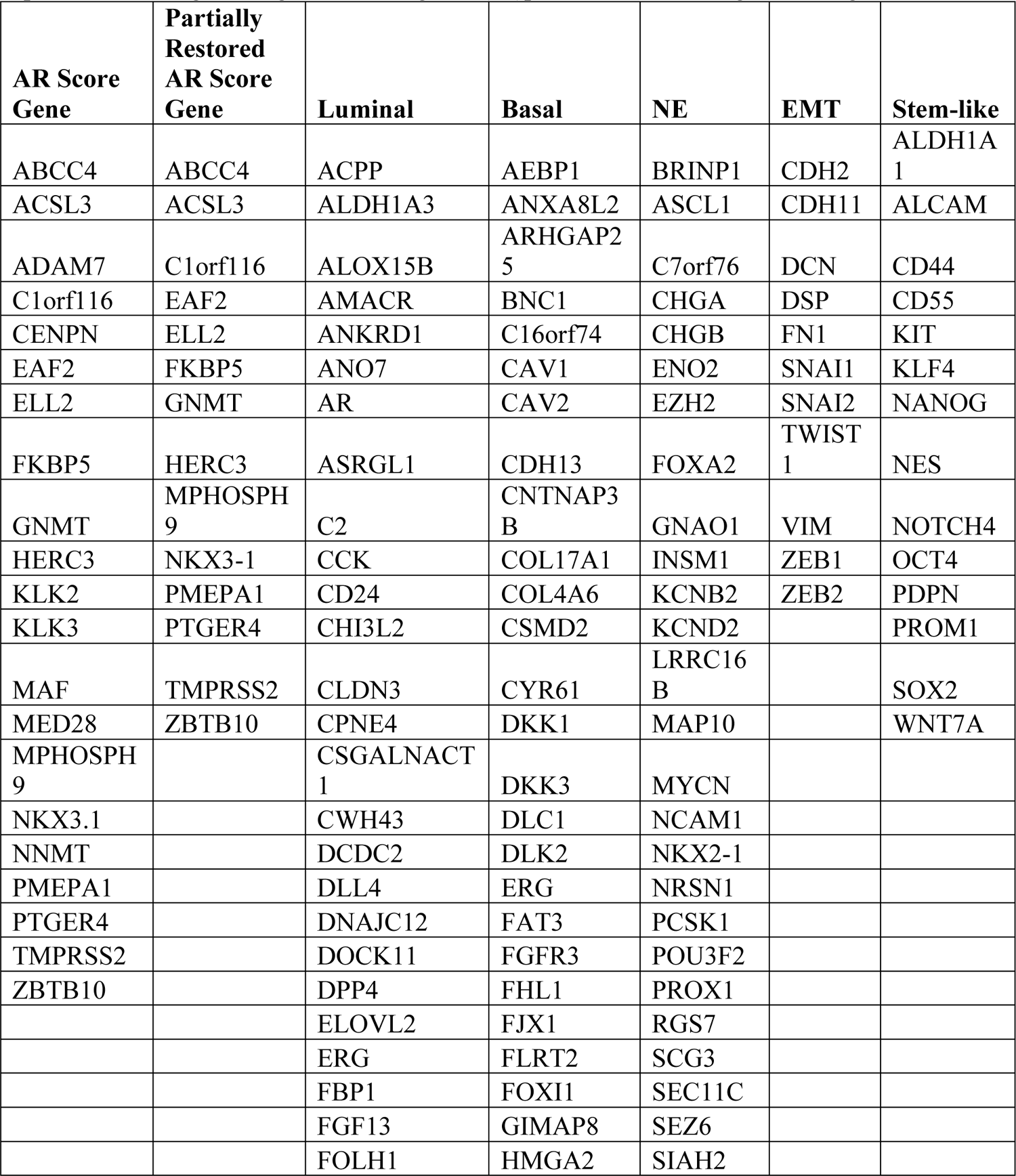
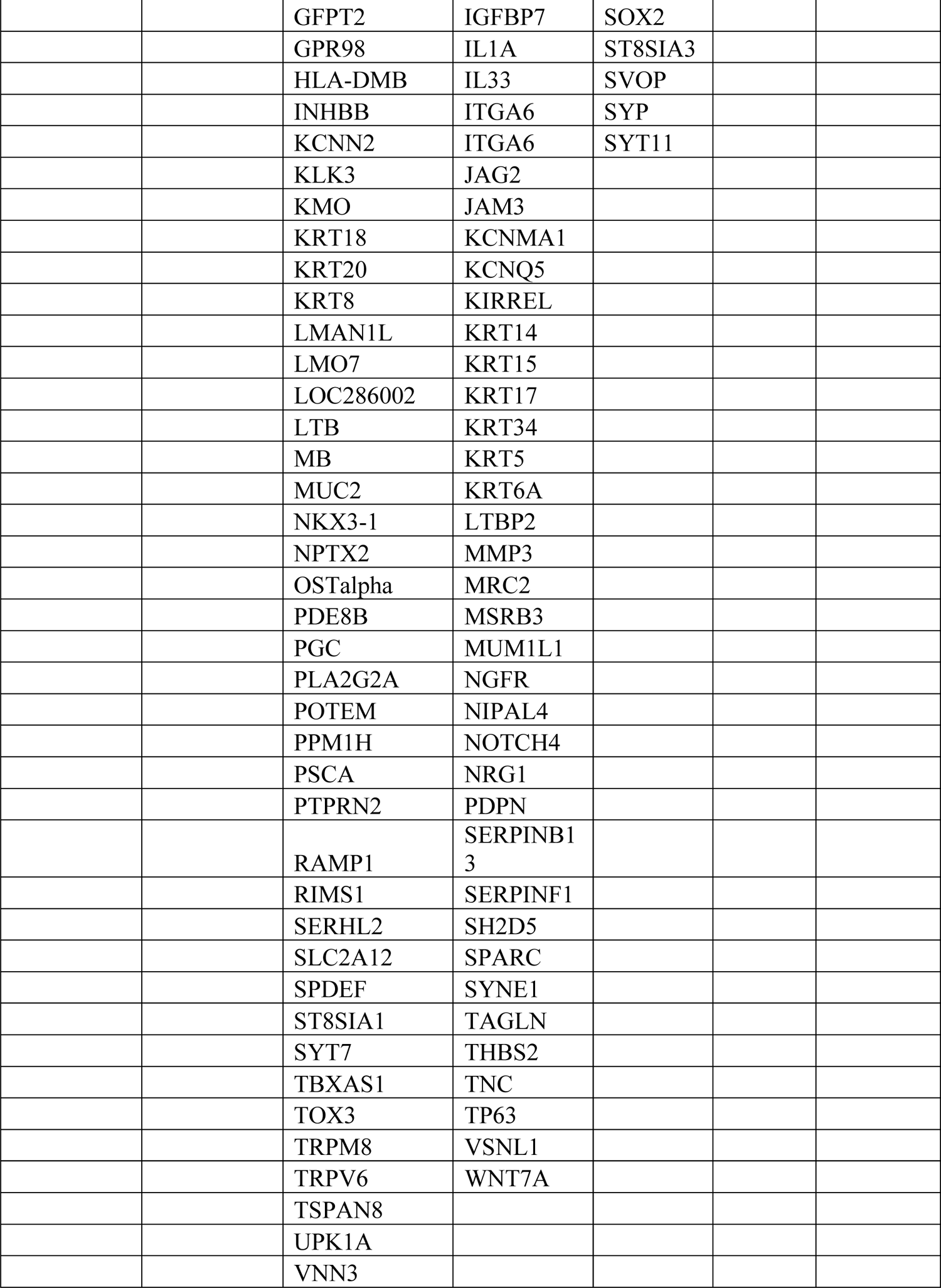
AR Score and lineage specific signatures gene lists. The AR score gene signature was adapted from Hieronymus et al(*48*), luminal, basal and NE gene signatures were defined by combining the signature genes from(*10, 11, 46, 47*). EMT and stem-like gene signature were adapted from the signature genes of Dong et al(*47*)plus canonical lineage marker genes.

## References

1. T. M. Beer, A. J. Armstrong, D. E. Rathkopf, Y. Loriot, C. N. Sternberg, C. S. Higano, P. Iversen, S. Bhattacharya, J. Carles, S. Chowdhury, I. D. Davis, J. S. de Bono, C. P. Evans, K. Fizazi, A. M. Joshua, C.-S. Kim, G. Kimura, P. Mainwaring, H. Mansbach, K. Miller, S. B. Noonberg, F. Perabo, D. Phung, F. Saad, H. I. Scher, M.-E. Taplin, P. M. Venner, B. Tombal, P. Investigators, Enzalutamide in Metastatic Prostate Cancer before Chemotherapy. New Engl J Medicine. 371, 424–433 (2014).

2. C. J. Ryan, M. R. Smith, J. S. de Bono, A. Molina, C. J. Logothetis, P. de Souza, K. Fizazi, P. Mainwaring, J. M. Piulats, S. Ng, J. Carles, P. F. A. Mulders, E. Basch, E. J. Small, F. Saad, D. Schrijvers, H. V. Poppel, S. D. Mukherjee, H. Suttmann, W. R. Gerritsen, T. W. Flaig, D. J. George, E. Y. Yu, E. Efstathiou, A. Pantuck, E. Winquist, C. S. Higano, M.-E. Taplin, Y. Park, T. Kheoh, T. Griffin, H. I. Scher, D. E. Rathkopf, C.-A.-302 Investigators, Abiraterone in Metastatic Prostate Cancer without Previous Chemotherapy. New Engl J Medicine. 368, 138–148 (2013).

3. M. R. Smith, F. Saad, S. Chowdhury, S. Oudard, B. A. Hadaschik, J. N. Graff, D. Olmos, P. N. Mainwaring, J. Y. Lee, H. Uemura, A. Lopez-Gitlitz, G. C. Trudel, B. M. Espina, Y. Shu, Y. C. Park, W. R. Rackoff, M. K. Yu, E. J. Small, Apalutamide Treatment and Metastasis-free Survival in Prostate Cancer. New Engl J Medicine. 378, 1408–1418 (2018).

4. P. A. Watson, V. K. Arora, C. L. Sawyers, Nat Rev Cancer, in press, doi:10.1038/nrc4016.

5. V. K. Arora, E. Schenkein, R. Murali, S. K. Subudhi, J. Wongvipat, M. D. Balbas, N. Shah, L. Cai, E. Efstathiou, C. Logothetis, D. Zheng, C. L. Sawyers, Cell, in press, doi:10.1016/j.cell.2013.11.012.

6. M. Isikbay, K. Otto, S. Kregel, J. Kach, Y. Cai, D. J. V. Griend, S. D. Conzen, R. Z. Szmulewitz, Glucocorticoid Receptor Activity Contributes to Resistance to Androgen-Targeted Therapy in Prostate Cancer. Hormones Cancer. 5, 72–89 (2014).

7. Z. Zhang, C. Zhou, X. Li, S. D. Barnes, S. Deng, E. Hoover, C.-C. Chen, Y. S. Lee, Y. Zhang, C. Wang, L. A. Metang, C. Wu, C. R. Tirado, N. A. Johnson, J. Wongvipat, K. Navrazhina, Z. Cao, D. Choi, C.-H. Huang, E. Linton, X. Chen, Y. Liang, C. E. Mason, E. de Stanchina, W. Abida, A. Lujambio, S. Li, S. W. Lowe, J. T. Mendell, V. S. Malladi, C. L. Sawyers, P. Mu, Loss of CHD1 Promotes Heterogeneous Mechanisms of Resistance to AR-Targeted Therapy via Chromatin Dysregulation. Cancer Cell (2020), doi:10.1016/j.ccell.2020.03.001.

8. H. Beltran, A. Hruszkewycz, H. I. Scher, J. Hildesheim, J. Isaacs, E. Y. Yu, K. Kelly, D. Lin, A. Dicker, J. Arnold, T. Hecht, M. Wicha, R. Sears, D. Rowley, R. White, J. L. Gulley, J. Lee, M. D. Meco, E. J. Small, M. Shen, K. Knudsen, D. W. Goodrich, T. Lotan, A. Zoubeidi, C. L. Sawyers, C. M. Rudin, M. Loda, T. Thompson, M. A. Rubin, A. Tawab-Amiri, W. Dahut, P. S. Nelson, The Role of Lineage Plasticity in Prostate Cancer Therapy Resistance. Clin Cancer Res. 25, 6916– 6924 (2019).

9. S. Y. Ku, S. Rosario, Y. Wang, P. Mu, M. Seshadri, Z. W. Goodrich, M. M. Goodrich, D. P. Labbé, E. C. Gomez, J. Wang, H. W. Long, B. Xu, M. Brown, M. Loda, C. L. Sawyers, L. Ellis, D. W. Goodrich, Science, in press, doi:10.1126/science.aah4199.

10. P. Mu, Z. Zhang, M. Benelli, W. R. Karthaus, E. Hoover, C.-C. Chen, J. Wongvipat, S. Y. Ku, D. Gao, Z. Cao, N. Shah, E. J. Adams, W. Abida, P. A. Watson, D. Prandi, C.-H. Huang, E. de Stanchina, S. W. Lowe, L. Ellis, H. Beltran, M. A. Rubin, D. W. Goodrich, F. Demichelis, C. L. Sawyers, Science, in press, doi:10.1126/science.aah4307.

11. H. Beltran, D. Prandi, J.-M. Mosquera, M. Benelli, L. Puca, J. Cyrta, C. Marotz, E. Giannopoulou, B. V. S. K. Chakravarthi, S. Varambally, S. A. Tomlins, D. M. Nanus, S. T. Tagawa, E. M. V. Allen, O. Elemento, A. Sboner, L. A. Garraway, M. A. Rubin, F. Demichelis, Nat Med, in press, doi:10.1038/nm.4045.

12. E. Dardenne, H. Beltran, M. Benelli, K. Gayvert, A. Berger, L. Puca, J. Cyrta, A. Sboner, Z. Noorzad, T. MacDonald, C. Cheung, K. S. Yuen, D. Gao, Y. Chen, M. Eilers, J.-M. Mosquera, B. D. Robinson, O. Elemento, M. A. Rubin, F. Demichelis, D. S. Rickman, N-Myc Induces an EZH2-Mediated Transcriptional Program Driving Neuroendocrine Prostate Cancer. Cancer Cell. 30, 563–577 (2016).

13. J. L. Bishop, D. Thaper, S. Vahid, A. Davies, K. Ketola, H. Kuruma, R. Jama, K. M. Nip, A. Angeles, F. Johnson, A. W. Wyatt, L. Fazli, M. E. Gleave, D. Lin, M. A. Rubin, C. C. Collins, Y. Wang, H. Beltran, A. Zoubeidi, The Master Neural Transcription Factor BRN2 Is an Androgen Receptor–Suppressed Driver of Neuroendocrine Differentiation in Prostate Cancer. Cancer Discov. 7, 54–71 (2017).

14. J. Cyrta, A. Augspach, M. R. de Filippo, D. Prandi, P. Thienger, M. Benelli, V. Cooley, R. Bareja, D. Wilkes, S.-S. Chae, P. Cavaliere, N. Dephoure, A.-C. Uldry, S. B. Lagache, S. Cohen, M. Jaquet, L. P. Brandt, M. Alshalalfa, A. Sboner, F. Feng, S. Wang, H. Beltran, T. Lotan, M. Spahn, M. K. Julio, Y. Chen, K. V. Ballman, F. Demichelis, S. Piscuoglio, M. A. Rubin, Biorxiv, in press, doi:10.1101/2020.03.06.949131.

15. M. Zou, R. Toivanen, A. Mitrofanova, N. Floc’h, S. Hayati, Y. Sun, C. L. Magnen, D. Chester, E. A. Mostaghel, A. Califano, M. A. Rubin, M. M. Shen, C. Abate-Shen, Transdifferentiation as a Mechanism of Treatment Resistance in a Mouse Model of Castration-resistant Prostate Cancer. Cancer Discov. 7, 736–749 (2017).

16. X. Zhang, I. M. Coleman, L. G. Brown, L. D. True, L. Kollath, J. M. Lucas, H.-M. Lam, R. Dumpit, E. Corey, L. Chéry, B. Lakely, C. S. Higano, B. Montgomery, M. Roudier, P. H. Lange, P. S. Nelson, R. L. Vessella, C. Morrissey, SRRM4 Expression and the Loss of REST Activity May Promote the Emergence of the Neuroendocrine Phenotype in Castration-Resistant Prostate Cancer. Am Assoc Cancer Res. 21, 4698–4708 (2015).

17. E. J. Adams, W. R. Karthaus, E. Hoover, D. Liu, A. Gruet, Z. Zhang, H. Cho, R. DiLoreto, S. Chhangawala, Y. Liu, P. A. Watson, E. Davicioni, A. Sboner, C. E. Barbieri, R. Bose, C. S. Leslie, C. L. Sawyers, FOXA1 mutations alter pioneering activity, differentiation and prostate cancer phenotypes. Nature. 571, 408–412 (2019).

18. A. Davies, S. Nouruzi, D. Ganguli, T. Namekawa, D. Thaper, S. Linder, F. Karaoğlanoğlu, M. E. Omur, S. Kim, M. Kobelev, S. Kumar, O. Sivak, C. Bostock, J. Bishop, M. Hoogstraat, A. Talal, S. Stelloo, H. van der Poel, A. M. Bergman, M. Ahmed, L. Fazli, H. Huang, W. Tilley, D. Goodrich, F. Y. Feng, M. Gleave, H. H. He, F. Hach, W. Zwart, H. Beltran, L. Selth, A. Zoubeidi, An androgen receptor switch underlies lineage infidelity in treatment-resistant prostate cancer. Nat Cell Biol. 23, 1023–1034 (2021).

19. L. A. Garraway, H. R. Widlund, M. A. Rubin, G. Getz, A. J. Berger, S. Ramaswamy, R. Beroukhim, D. A. Milner, S. R. Granter, J. Du, C. Lee, S. N. Wagner, C. Li, T. R. Golub, D. L. Rimm, M. L. Meyerson, D. E. Fisher, W. R. Sellers, Integrative genomic analyses identify MITF as a lineage survival oncogene amplified in malignant melanoma. Nature. 436, 117 (2005).

20. J. W. Park, J. K. Lee, K. M. Sheu, L. Wang, N. G. Balanis, K. Nguyen, B. A. Smith, C. Cheng, B. L. Tsai, D. Cheng, J. Huang, S. K. Kurdistani, T. G. Graeber, O. N. Witte, Reprogramming normal human epithelial tissues to a common, lethal neuroendocrine cancer lineage. Science. 362, 91–95 (2018).

21. L. V. Sequist, B. A. Waltman, D. Dias-Santagata, S. Digumarthy, A. B. Turke, P. Fidias, K. Bergethon, A. T. Shaw, S. Gettinger, A. K. Cosper, S. Akhavanfard, R. S. Heist, J. Temel, J. G. Christensen, J. C. Wain, T. J. Lynch, K. Vernovsky, E. J. Mark, M. Lanuti, A. J. Iafrate, M. Mino-Kenudson, J. A. Engelman, Sci Transl Med, in press, doi:10.1126/scitranslmed.3002003.

22. G. Xu, S. Chhangawala, E. Cocco, P. Razavi, Y. Cai, J. E. Otto, L. Ferrando, P. Selenica, E. Ladewig, C. Chan, A. D. C. Paula, M. Witkin, Y. Cheng, J. Park, C. Serna-Tamayo, H. Zhao, F. Wu, M. Sallaku, X. Qu, A. Zhao, C. K. Collings, A. R. D’Avino, K. Jhaveri, R. Koche, R. L. Levine, J. S. Reis-Filho, C. Kadoch, M. Scaltriti, C. S. Leslie, J. Baselga, E. Toska, ARID1A determines luminal identity and therapeutic response in estrogen-receptor-positive breast cancer. Nat Genet. 52, 198–207 (2020).

23. S. J. Thomas, J. A. Snowden, M. P. Zeidler, S. J. Danson, The role of JAK/STAT signalling in the pathogenesis, prognosis and treatment of solid tumours. Brit J Cancer. 113, 365–371 (2015).

24. S. R. Singh, X. Chen, S. X. Hou, JAK/STAT signaling regulates tissue outgrowth and male germline stem cell fate in Drosophila. Cell Res. 15, 1–5 (2005).

25. K. Beebe, W.-C. Lee, C. A. Micchelli, JAK/STAT signaling coordinates stem cell proliferation and multilineage differentiation in the Drosophila intestinal stem cell lineage. Dev Biol. 338, 28– 37 (2010).

26. T. Mirtti, B. E. Leiby, J. Abdulghani, E. Aaltonen, M. Pavela, A. Mamtani, K. Alanen, L. Egevad, T. Granfors, A. Josefsson, P. Stattin, A. Bergh, M. T. Nevalainen, Nuclear Stat5a/b predicts early recurrence and prostate cancer–specific death in patients treated by radical prostatectomy. Hum Pathol. 44, 310–319 (2013).

27. K. L. Owen, N. K. Brockwell, B. S. Parker, JAK-STAT Signaling: A Double-Edged Sword of Immune Regulation and Cancer Progression. Cancers. 11, 2002 (2019).

28. K. Meissl, S. Macho-Maschler, M. Müller, B. Strobl, The good and the bad faces of STAT1 in solid tumours. Cytokine. 89, 12–20 (2017).

29. M. T. Spiotto, T. D. K. Chung, STAT3 mediates IL-6-induced neuroendocrine differentiation in prostate cancer cells. Prostate. 42, 186–195 (2000).

30. Y. Zhu, C. Liu, Y. Cui, N. Nadiminty, W. Lou, A. C. Gao, Interleukin-6 induces neuroendocrine differentiation (NED) through suppression of RE-1 silencing transcription factor (REST). Prostate. 74, 1086–1094 (2014).

31. J. Kim, R. M. Adam, K. R. Solomon, M. R. Freeman, Involvement of Cholesterol-Rich Lipid Rafts in Interleukin-6-Induced Neuroendocrine Differentiation of LNCaP Prostate Cancer Cells. Endocrinology. 145, 613–619 (2004).

32. S. O. Lee, J. Y. Chun, N. Nadiminty, W. Lou, A. C. Gao, Interleukin-6 undergoes transition from growth inhibitor associated with neuroendocrine differentiation to stimulator accompanied by androgen receptor activation during LNCaP prostate cancer cell progression. Prostate. 67, 764– 773 (2007).

33. S. O. Lee, W. Lou, C. S. Johnson, D. L. Trump, A. C. Gao, Interleukin-6 protects LNCaP cells from apoptosis induced by androgen deprivation through the Stat3 pathway. Prostate. 60, 178– 186 (2004).

34. S. O. Lee, W. Lou, M. Hou, F. de Miguel, L. Gerber, A. C. Gao, Interleukin-6 promotes androgen-independent growth in LNCaP human prostate cancer cells. Clin Cancer Res Official J Am Assoc Cancer Res. 9, 370–6 (2003).

35. C. Liu, W. Lou, C. Armstrong, Y. Zhu, C. P. Evans, A. C. Gao, Niclosamide suppresses cell migration and invasion in enzalutamide resistant prostate cancer cells via Stat3-AR axis inhibition. Prostate. 75, 1341–1353 (2015).

36. C. Liu, Y. Zhu, W. Lou, Y. Cui, C. P. Evans, A. C. Gao, Inhibition of constitutively active Stat3 reverses enzalutamide resistance in LNCaP derivative prostate cancer cells. Prostate. 74, 201–209 (2014).

37. K. H. Cho, K. J. Jeong, S. C. Shin, J. Kang, C. G. Park, H. Y. Lee, STAT3 mediates TGF-β1-induced TWIST1 expression and prostate cancer invasion. Cancer Lett. 336, 167–173 (2013).

38. A. Rojas, G. Liu, I. Coleman, P. S. Nelson, M. Zhang, R. Dash, P. B. Fisher, S. R. Plymate, J. D. Wu, IL-6 promotes prostate tumorigenesis and progression through autocrine cross-activation of IGF-IR. Oncogene. 30, 2345–2355 (2011).

39. M. Sun, C. Liu, N. Nadiminty, W. Lou, Y. Zhu, J. Yang, C. P. Evans, Q. Zhou, A. C. Gao, Inhibition of Stat3 activation by sanguinarine suppresses prostate cancer cell growth and invasion. Prostate. 72, 82–89 (2012).

40. F. DeMiguel, S. O. Lee, W. Lou, X. Xiao, B. R. Pflug, J. B. Nelson, A. C. Gao, Stat3 enhances the growth of LNCaP human prostate cancer cells in intact and castrated male nude mice. Prostate. 52, 123–129 (2002).

41. S. O. Lee, W. Lou, K. M. Qureshi, F. Mehraein-Ghomi, D. L. Trump, A. C. Gao, RNA interference targeting Stat3 inhibits growth and induces apoptosis of human prostate cancer cells. Prostate. 60, 303–309 (2004).

42. W. Abida, J. Cyrta, G. Heller, D. Prandi, J. Armenia, I. Coleman, M. Cieslik, M. Benelli, D. Robinson, E. M. V. Allen, A. Sboner, T. Fedrizzi, J. M. Mosquera, B. D. Robinson, N. D. Sarkar, L. P. Kunju, S. Tomlins, Y. M. Wu, D. N. Rodrigues, M. Loda, A. Gopalan, V. E. Reuter, C. C. Pritchard, J. Mateo, D. Bianchini, S. Miranda, S. Carreira, P. Rescigno, J. Filipenko, J. Vinson, R. B. Montgomery, H. Beltran, E. I. Heath, H. I. Scher, P. W. Kantoff, M.-E. Taplin, N. Schultz, J. S. deBono, F. Demichelis, P. S. Nelson, M. A. Rubin, A. M. Chinnaiyan, C. L. Sawyers, Genomic correlates of clinical outcome in advanced prostate cancer. Proc National Acad Sci, 201902651 (2019).

43. C. G. A. R. Network, Cell, in press, doi:10.1016/j.cell.2015.10.025.

44. W. Hessenkemper, J. Roediger, S. Bartsch, A. B. Houtsmuller, M. E. van Royen, I. Petersen, M.-O. Grimm, A. Baniahmad, A Natural Androgen Receptor Antagonist Induces Cellular Senescence in Prostate Cancer Cells. Mol Endocrinol. 28, 1831–1840 (2014).

45. L. McInnes, J. Healy, J. Melville, UMAP: Uniform Manifold Approximation and Projection for Dimension Reduction. *Arxiv* (2018).

46. D. Zhang, D. Park, Y. Zhong, Y. Lu, K. Rycaj, S. Gong, X. Chen, X. Liu, H.-P. Chao, P. Whitney, T. Calhoun-Davis, Y. Takata, J. Shen, V. R. Iyer, D. G. Tang, Stem cell and neurogenic gene-expression profiles link prostate basal cells to aggressive prostate cancer. Nat Commun. 7, 10798 (2016).

47. B. Dong, J. Miao, Y. Wang, W. Luo, Z. Ji, H. Lai, M. Zhang, X. Cheng, J. Wang, Y. Fang, H. H. Zhu, C. W. Chua, L. Fan, Y. Zhu, J. Pan, J. Wang, W. Xue, W.-Q. Gao, Single-cell analysis supports a luminal-neuroendocrine transdifferentiation in human prostate cancer. Commun Biology. 3, 778 (2020).

48. H. Hieronymus, J. Lamb, K. N. Ross, X. P. Peng, C. Clement, A. Rodina, M. Nieto, J. Du, K. Stegmaier, S. M. Raj, K. N. Maloney, J. Clardy, W. C. Hahn, G. Chiosis, T. R. Golub, Gene expression signature-based chemical genomic prediction identifies a novel class of HSP90 pathway modulators. Cancer Cell. 10, 321–330 (2006).

49. M. D. Nyquist, A. Corella, I. Coleman, N. D. Sarkar, A. Kaipainen, G. Ha, R. Gulati, L. Ang, P. Chatterjee, J. Lucas, C. Pritchard, G. Risbridger, J. Isaacs, B. Montgomery, C. Morrissey, E. Corey, P. S. Nelson, Combined TP53 and RB1 Loss Promotes Prostate Cancer Resistance to a Spectrum of Therapeutics and Confers Vulnerability to Replication Stress. Cell Reports. 31, 107669 (2020).

50. I. Dagogo-Jack, A. T. Shaw, Tumour heterogeneity and resistance to cancer therapies. Nat Rev Clin Oncol. 15, 81–94 (2017).

51. C. Yao, L. Su, J. Shan, C. Zhu, L. Liu, C. Liu, Y. Xu, Z. Yang, X. Bian, J. Shao, J. Li, M. Lai, J. Shen, C. Qian, IGF/STAT3/NANOG/Slug Signaling Axis Simultaneously Controls Epithelial-Mesenchymal Transition and Stemness Maintenance in Colorectal Cancer. Stem Cells. 34, 820– 831 (2016).

52. A. Yadav, B. Kumar, J. Datta, T. N. Teknos, P. Kumar, IL-6 Promotes Head and Neck Tumor Metastasis by Inducing Epithelial–Mesenchymal Transition via the JAK-STAT3-SNAIL Signaling Pathway. Mol Cancer Res. 9, 1658–1667 (2011).

53. H. Xiong, J. Hong, W. Du, Y. Lin, L. Ren, Y. Wang, W. Su, J. Wang, Y. Cui, Z. Wang, J.-Y. Fang, Roles of STAT3 and ZEB1 Proteins in E-cadherin Down-regulation and Human Colorectal Cancer Epithelial-Mesenchymal Transition*. J Biol Chem. 287, 5819–5832 (2012).

54. C. Gong, J. Shen, Z. Fang, L. Qiao, R. Feng, X. Lin, S. Li, Abnormally expressed JunB transactivated by IL-6/STAT3 signaling promotes uveal melanoma aggressiveness via epithelial– mesenchymal transition. Bioscience Rep. 38 (2018), doi:10.1042/bsr20180532.

55. W. Malilas, S. S. Koh, S. Kim, R. Srisuttee, I.-R. Cho, J. Moon, H.-S. Yoo, S. Oh, R. N. Johnston, Y.-H. Chung, Cancer upregulated gene 2, a novel oncogene, enhances migration and drug resistance of colon cancer cells via STAT1 activation. Int J Oncol. 43, 1111– 1116 (2013).

56. N. N. Khodarev, M. Beckett, E. Labay, T. Darga, B. Roizman, R. R. Weichselbaum, STAT1 is overexpressed in tumors selected for radioresistance and confers protection from radiation in transduced sensitive cells. P Natl Acad Sci Usa. 101, 1714–1719 (2004).

57. G. S. Wong, J.-S. Lee, Y.-Y. Park, A. J. Klein-Szanto, T. J. Waldron, E. Cukierman, M. Herlyn, P. Gimotty, H. Nakagawa, A. K. Rustgi, Periostin cooperates with mutant p53 to mediate invasion through the induction of STAT1 signaling in the esophageal tumor microenvironment. Oncogenesis. 2, e59–e59 (2013).

58. C. Greenwood, G. Metodieva, K. Al-Janabi, B. Lausen, L. Alldridge, L. Leng, R. Bucala, N. Fernandez, M. V. Metodiev, Stat1 and CD74 overexpression is co-dependent and linked to increased invasion and lymph node metastasis in triple-negative breast cancer. J Proteomics. 75, 3031–3040 (2012).

59. M. Offin, J. M. Chan, M. Tenet, H. A. Rizvi, R. Shen, G. J. Riely, N. Rekhtman, Y. Daneshbod, A. Quintanal-Villalonga, A. Penson, M. D. Hellmann, M. E. Arcila, M. Ladanyi, D. Pe’er, M. G. Kris, C. M. Rudin, H. A. Yu, Concurrent RB1 and TP53 Alterations Define a Subset of EGFR-Mutant Lung Cancers at risk for Histologic Transformation and Inferior Clinical Outcomes. J Thorac Oncol. 14, 1784–1793 (2019).

60. M. Girardot, C. Pecquet, I. Chachoua, J. V. Hees, S. Guibert, A. Ferrant, L. Knoops, E. J. Baxter, P. A. Beer, S. Giraudier, R. Moriggl, W. Vainchenker, A. R. Green, S. N. Constantinescu, Persistent STAT5 activation in myeloid neoplasms recruits p53 into gene regulation. Oncogene. 34, 1323–1332 (2015).

61. E. Colombo, J.-C. Marine, D. Danovi, B. Falini, P. G. Pelicci, Nucleophosmin regulates the stability and transcriptional activity of p53. Nat Cell Biol. 4, 529–533 (2002).

62. H. J. Park, J. Li, R. Hannah, S. Biddie, A. I. Leal-Cervantes, K. Kirschner, D. F. S. Cruz, V. Sexl, B. Göttgens, A. R. Green, Cytokine-induced megakaryocytic differentiation is regulated by genome-wide loss of a uSTAT transcriptional program. Embo J. 35, 580–594 (2016).

63. C. D. Chen, D. S. Welsbie, C. Tran, S. H. Baek, R. Chen, R. Vessella, M. G. Rosenfeld, C. L. Sawyers, Nat Med, in press, doi:10.1038/nm972.

64. K. A. Klein, R. E. Reiter, J. Redula, H. Moradi, X. L. Zhu, A. R. Brothman, D. J. Lamb, M. Marcelli, A. Belldegrun, O. N. Witte, C. L. Sawyers, Progression of metastatic human prostate cancer to androgen independence in immunodeficient SCID mice. Nat Med. 3, 402–408 (1997).

65. W. R. Karthaus, P. J. Iaquinta, J. Drost, A. Gracanin, R. van Boxtel, J. Wongvipat, C. M. Dowling, D. Gao, H. Begthel, N. Sachs, R. G. J. Vries, E. Cuppen, Y. Chen, C. L. Sawyers, H. C. Clevers, Cell, in press, doi:10.1016/j.cell.2014.08.017.

66. D. Gao, I. Vela, A. Sboner, P. J. Iaquinta, W. R. Karthaus, A. Gopalan, C. Dowling, J. N. Wanjala, E. A. Undvall, V. K. Arora, J. Wongvipat, M. Kossai, S. Ramazanoglu, L. P. Barboza, W. Di, Z. Cao, Q. F. Zhang, I. Sirota, L. Ran, T. Y. MacDonald, H. Beltran, J.-M. Mosquera, K. A. Touijer, P. T. Scardino, V. P. Laudone, K. R. Curtis, D. E. Rathkopf, M. J. Morris, D. C. Danila, S. F. Slovin, S. B. Solomon, J. A. Eastham, P. Chi, B. Carver, M. A. Rubin, H. I. Scher, H. Clevers, C. L. Sawyers, Y. Chen, Cell, in press, doi:10.1016/j.cell.2014.08.016.

67. D. B. Wheeler, R. Zoncu, D. E. Root, D. M. Sabatini, C. L. Sawyers, Science, in press, doi:10.1126/science.aaa4903.

68. F. A. Wolf, P. Angerer, F. J. Theis, SCANPY: large-scale single-cell gene expression data analysis. Genome Biol. 19, 15 (2018).

69. V. A. Traag, L. Waltman, N. J. van Eck, From Louvain to Leiden: guaranteeing well-connected communities. Sci Rep-uk. 9, 5233 (2019).

